# White Matter Signals Reflect Information Transmission Between Brain Regions During Seizures

**DOI:** 10.1101/2021.09.15.460549

**Authors:** Andrew Y. Revell, Alexander B. Silva, Dhanya Mahesh, Lena Armstrong, T. Campbell Arnold, John M. Bernabei, Ezequiel Gleichgerrcht, Leonardo Bonilha, Joel M. Stein, Sandhitsu R. Das, Russell T. Shinohara, Dani S. Bassett, Brian Litt, Kathryn A. Davis

## Abstract

White matter supports critical brain functions such as learning and memory, modulates the distribution of action potentials, and transmits neural information between brain regions. Notably, neuronal cell bodies exist in deeper white matter tissue, neurotransmitter vesicles are released directly in white matter, and white matter blood-oxygenation level dependent (BOLD) signals are detectable across a range of different tasks—all appearing to reflect a dynamic, active tissue from which recorded signals can reveal meaningful information about the brain. Yet, the signals within white matter have largely been ignored. Here, we elucidate the properties of white matter signals using intracranial EEG in a bipolar montage. We show that such signals capture the communication between brain regions and differentiate pathophysiologies of epilepsy. In direct contradiction to past assumptions that white matter functional signals provide little value, we show that white matter recordings can elucidate brain function and pathophysiology. Broadly, white matter functional recordings acquired through implantable devices may provide a wealth of currently untapped knowledge about the neurobiology of disease.

## Introduction

Stereoelectroencephalography (SEEG) provides vital neural information in medically refractory epilepsy patients to aid in localization of the seizure onset zone (SOZ) for its eventual resection, ablation, or modulation (Fig. 1a)^1,2^. This implantation modality is now favored in clinical practice because it decreases morbidity and is better tolerated by patients^3^. It also provides coverage to deeper brain structures, capturing electrical activity in many tissue types, including white matter (Fig. 1b).

**Fig. 1.**
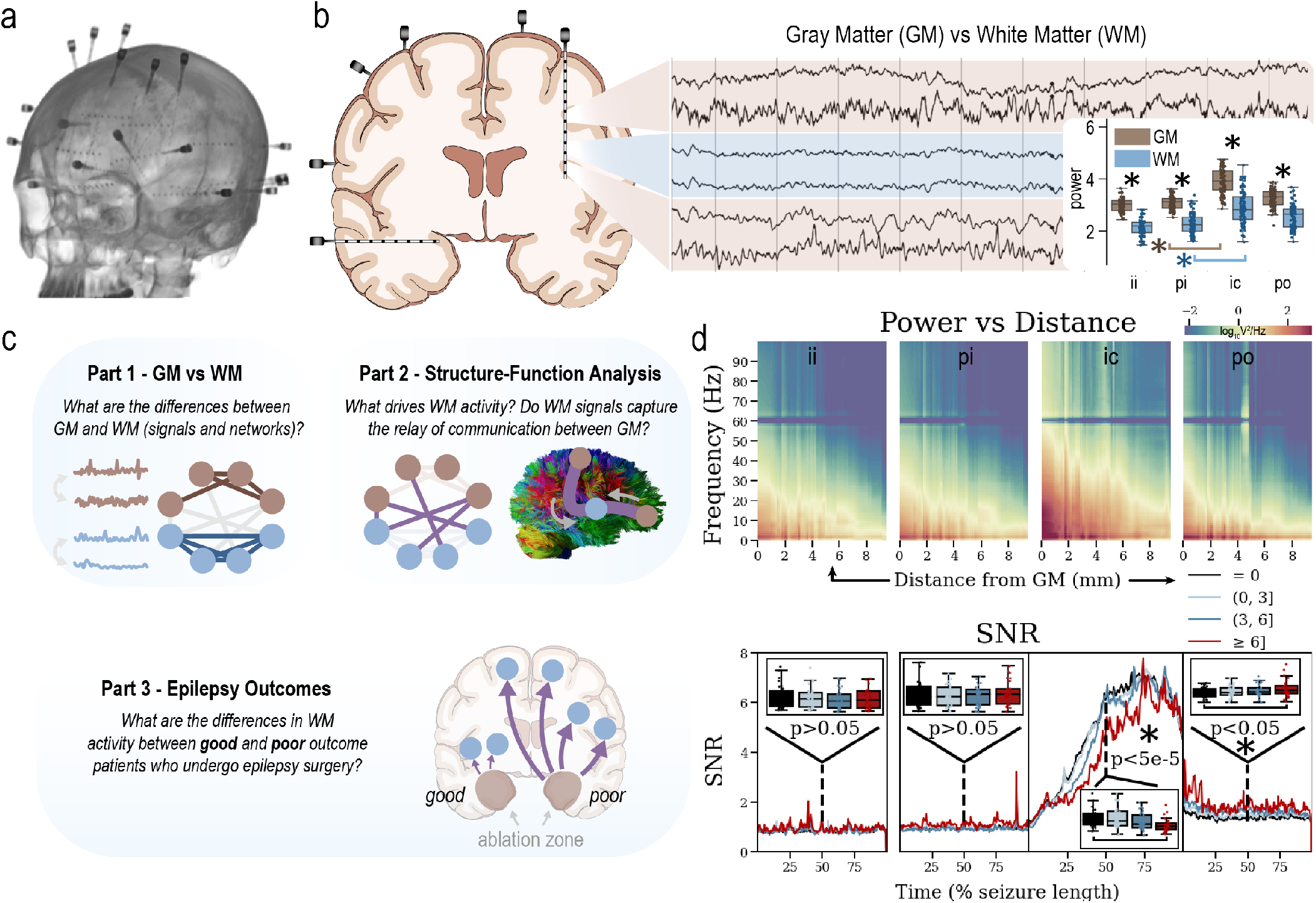
White matter recordings may elucidate brain function and pathophysiology. **a**, A CT scan of an epilepsy patient who underwent bilateral stereoelectroencephalography (SEEG) implantation. **b**, SEEG records from both gray matter (GM) and white matter (WM). Recordings can appear visually different along the depth of an SEEG electrode, which can be problematic for localizing the seizure onset zone^4^. A power analysis (inset) shows that WM recordings have lower power than GM recordings (interictal, preictal, ictal, postictal). **c**, Here, we investigate WM recordings in three parts: (1) assess differences between the signals and networks of GM and WM, (2) understand what drives WM signals, and (3) differentiate good and poor outcome epilepsy patients with WM recordings. **d**, Power in WM decreases farther from GM tissue (top). WM contacts ≥ 6mm from GM tissue have lower power during seizures compared to baseline (i.e. interictal power, *p* < 5*x*10^−5^) than contacts in GM tissue (bottom). However, power remains elevated *postictally* in WM, while power in GM tissue returns to baseline (*p* < 0.05). **SNR**, signal-to-noise ratio; **ii**, interictal; **pi**, preictal; **ic**, ictal; **po**, postictal.

Approximately half of SEEG contacts localize to white matter^5^ (Fig. S7); however, clinicians in many epilepsy centers typically exclude white matter contacts when creating intracranial EEG (iEEG) montages to interpret, diagnose, and localize the SOZ. White matter contacts also tend to be excluded from electrical stimulation mapping^6^. It is commonly assumed these recordings provide less clinically relevant information or they capture redundant information from the volume conduction of nearby gray matter regions^7,8^. Research studies^5,9–12^ utilizing SEEG also typically exclude white matter contacts; their signals may prove problematic when creating computational models sensitive to EEG features if white matter signals are functionally unique. Indeed, analyses of *interictal* recordings of select EEG features have shown differences in white matter signals^7,8,13,14^.

Do white matter SEEG recordings provide clinically relevant information? Do they capture information transmission along the white matter tracts connecting brain regions, or are they merely redundant signals from nearby gray matter tissue?

Here, we investigate white matter electrical recordings through a multi-pronged approach (Fig. 1c): (1) characterizing white matter and gray matter signal and functional network differences across distinct brain states, (2) testing the hypothesis^7^ that white matter signals capture the communication between gray matter regions using multimodal structural and functional data, and (3) demonstrating that white matter activity differs across outcomes of epilepsy patients who underwent surgery to remove the SOZ. We provide evidence highlighting the importance of white matter signals, which may harbor key information regarding neurobiology, disease, and diagnoses.

## Results

### The power of white matter iEEG signals is lower during seizures, but remains elevated postictally compared to baseline

A power analysis (Fig. 1b, inset) shows both gray matter (GM) and white matter (WM) recordings (≥2mm from GM) increase in broadband power from preictal to ictal states (*p* < 5*x*10^−5^). GM recordings also have higher power than WM recordings in all peri-ictal states (interictal, preictal, ictal, postictal, *p* < 5*x*10^−5^), replicating prior interictal studies at different institutions with different patient populations^7,8^. This observation confirms the hypothesis that WM recordings display different signal properties. We show that power across all frequency bands decreases in WM tissue as a function of distance from GM tissue (Fig. 1c). When comparing power *to baseline* (i.e. interictal) power, WM contacts ≥ 6mm from GM tissue have lower power during seizures (*p* < 5*x*10^−5^) than contacts in GM tissue. However, power remains elevated postictally in these WM recordings, while power in GM tissue returns to baseline (*p*< 0.05). This elevation in WM power postictally suggests that WM recordings are functionally distinct from GM recordings in epilepsy patients. This may also indicate that inter-regional communication, detected by contacts inserted in the white matter communication pathways themselves, is abnormal postictally, even when gray matter activity might return to baseline.

### White matter signals are more correlated with each other than gray matter signals

Next, we compared GM and WM functional connectivity (Fig. 2). We defined functional connectivity (FC) using high-*γ*-band cross correlation (100-128Hz) in a bipolar montage (see Methods on frequency definitions and montaging). This band along with bipolar montaging represents relatively local activity dynamics that are largely unaffected by volume conduction^17^. The band is particularly relevant to our study because a well-developed literature implicates high-frequency oscillations and *γ*-band activity as drivers of epileptic activity^18,19^. We also repeat all major plots in a common average reference (CAR) in Fig. S8 and different frequency bands in Fig. S3, Fig. S4 and Fig. S9 showing similar results.

**Fig. 2.**
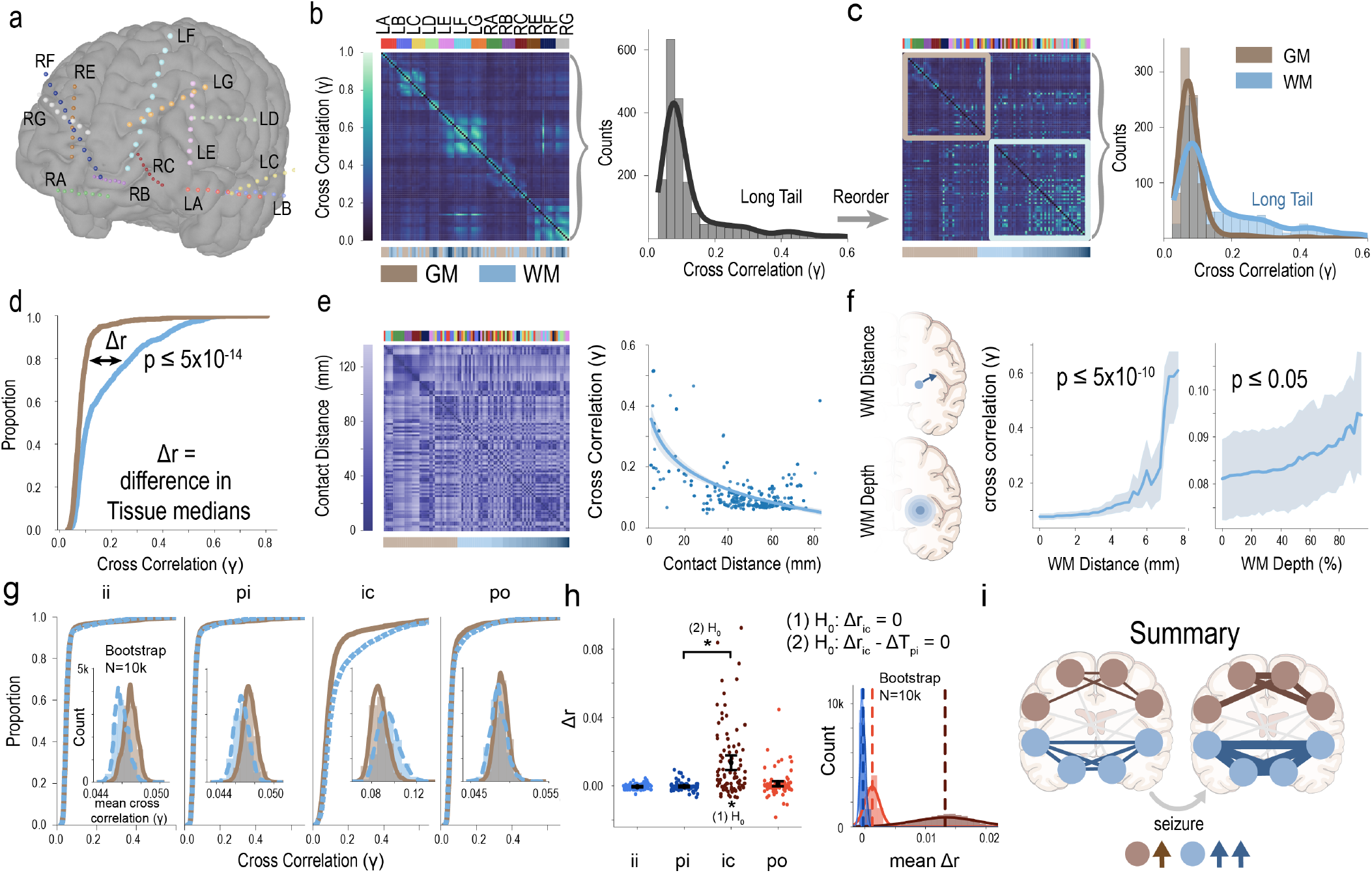
White matter (WM) correlations increase more than gray matter (GM) during seizures. **a**, SEEG with labeled electrodes. **b**, The high-*γ*-band cross correlation matrix ordered by electrode labels. Same color as in panel **a**. Bottom color bar indicates which contacts are GM versus WM. Distribution of FC (functional connectivity) values is plotted. **c**, Re-ordered adjacency matrix by WM distance from GM. FC distribution is separated by GM or WM values showing WM contributes to the tail. **d**, Empirical cumulative density function (ECDF) is plotted. Δ*r* is the difference in the median WM and GM FC values. **e**, Pairwise Euclidean distance between contacts (left). Contacts closer together have higher FC (right); therefore we used distance between contacts as a covariate in understanding the relationship between WM distance from GM and the increase in FC seen in panel **d. f**, WM contacts are determined by two definitions (Fig. S2). WM FC increases as a function of the respective WM definition after controlling for contact distance. Each WM definition is significantly related to FC. **g**, ECDF plots of WM and GM sub-networks during four states. The distribution of bootstrapped GM and WM FC values are inset. **h**, Δ*r* is plotted for each seizure and corresponding peri-ictal state (interictal, preictal, postictal). Error bars are 95% confidence intervals. Mean Δ*r* for each state of all bootstraps (*N* = 10, 000) are plotted. **i**, Summary of findings: WM correlations are higher than GM, and the increase in those correlations during seizures is greater in WM than in GM.

We localized SEEG contacts (Fig. 2a) to GM or WM (see the tissue localization subsection of the Methods), and plotted the distribution of all FC values (Fig. 2b) during the ictal state. We separated out tissue sub-networks (GM-only and WM-only) and observed that WM contributes to the tail of the distribution (Fig. 2c). Given that WM has lower power than GM and may record redundant noisy signals^13^, we initially hypothesized that WM FC would be *lower* than GM FC. We tested the null hypothesis that the two sub-network distributions of FC values were identical during the ictal period. Instead, we found that WM contacts have significantly *higher* FC values (*p*< 5*x*10^−14^, Fig. 2d) during the ictal state.

### Controlling for distances between contacts

We wanted to test if the higher FC values between WM contacts could be explained by the proximity between the WM contacts (i.e. the average distance between contacts inserted in each tissue type; not the inter-contact distance on an electrode, which is constant). It is reasonable to suspect that WM contacts are closer together than GM contacts because WM tissue is more centrally located in a given hemisphere, and some subcortical GM structures, such as the thalamus and putamen, are rarely implanted. However, contacts in medial GM structures (such as the cingulate gyrus or other limbic structures) are frequently targeted for bilateral implantation (thus these GM contacts are physically close). Indeed, contacts closer together have higher FC than contacts further away (Fig. 2e, Spearman Rank Correlation *r* = 0.31, *p*< 5*x*10^−10^), even in the high-*γ*-band after bipolar montaging. Thus, we used contact distance as a covariate. We fit an ordinary least squares regression model to predict FC as a function of the coefficients’ contact distance and either (1) distance from GM or (2) WM depth (see WM definition details in methods and Fig. S1 and S2). Briefly, we used two alternative definitions of WM because the limited number of studies analyzing interictal WM SEEG signals also used varying definitions, including distance to GM contacts and WM depth (where depth accounts for the surrounding tissue composition). A higher depth means the contact is deeply embedded within WM and has little surrounding GM tissue)^7,8^. Alternative definitions have shown similar results when comparing GM and WM signals in previous studies^8^, however, we still provide complementary analyses using both definitions here. We found that both distance from GM (*p* < 5*x*10^−10^) and WM depth (*p* < 0.05) were significantly related to FC (Fig. 2f). FC increases more in WM even after accounting for the distance between the WM contacts.

### White matter correlations increase more than gray matter signals during seizures

Next, we show the differences of FC between the tissue types across the four peri-ictal states (Fig. 2g-h). We show through bootstrapping (see the bootstrapping subsection of the Methods) that WM FC is slightly lower than GM FC during the interictal and preictal states, but increases dramatically more than GM during the ictal state (*p*< 0.05, Fig. 2g ECDF plots and inset histograms). The distributions of FC values in WM become similar to those in GM during the postictal state (*p* > 0.05). We test the null hypothesis that the difference in GM and WM median FC values during the ictal state is not zero (Δ*r*_*ic*_ ≠ 0, Fig. 2h). We found evidence that the median WM FC is higher than the median GM FC (Δ*r*_*ic*_ > 0; *p*< 0.005). Although both GM and WM FC increase from the preictal to the ictal states, we also show that the increase is larger for WM (*p* < 0.05). In summary (Fig. 2i), we show that WM FC is higher than GM FC, and that the increase in WM FC from the preictal to the ictal states is larger than in GM FC. We repeat our key results in broadband signals and show that the observed differences are even larger (Fig. S3). The values of Δ*r* for all bands and for cross correlation, coherence, and Pearson correlations are shown in Fig. S4 and support that these findings are not limited to specific bands or FC definitions.

### White matter sub-networks have different network properties than gray-matter sub-networks

As part of our multi-pronged approach, we compared GM and WM signals with a network analysis of the tissue sub-networks (Fig. 3). We measured density, characteristic path length, and transitivity of the full network which includes all contacts, the GM-only sub-network, and the WM-only sub-network. Density is a proportion of all possible connections that are actually present, characteristic path length is the average shortest distance between pairs of nodes, and transitivity is a measure of the tendency of the nodes to cluster together. We show that density and transitivity are both higher in WM-only sub-networks than in GM-only sub-networks during the ictal state (*p* < 0.05). Networks composed of only WM contacts are more clustered and have different network properties than networks composed of only GM contacts. This finding is crucial for informing research applying network models in SEEG. It remains to be seen whether the utility for network models excluding or including WM recordings affects clinical translation. However, we show here that functional network properties derived from data of both GM and WM tissue are affected.

**Fig. 3.**
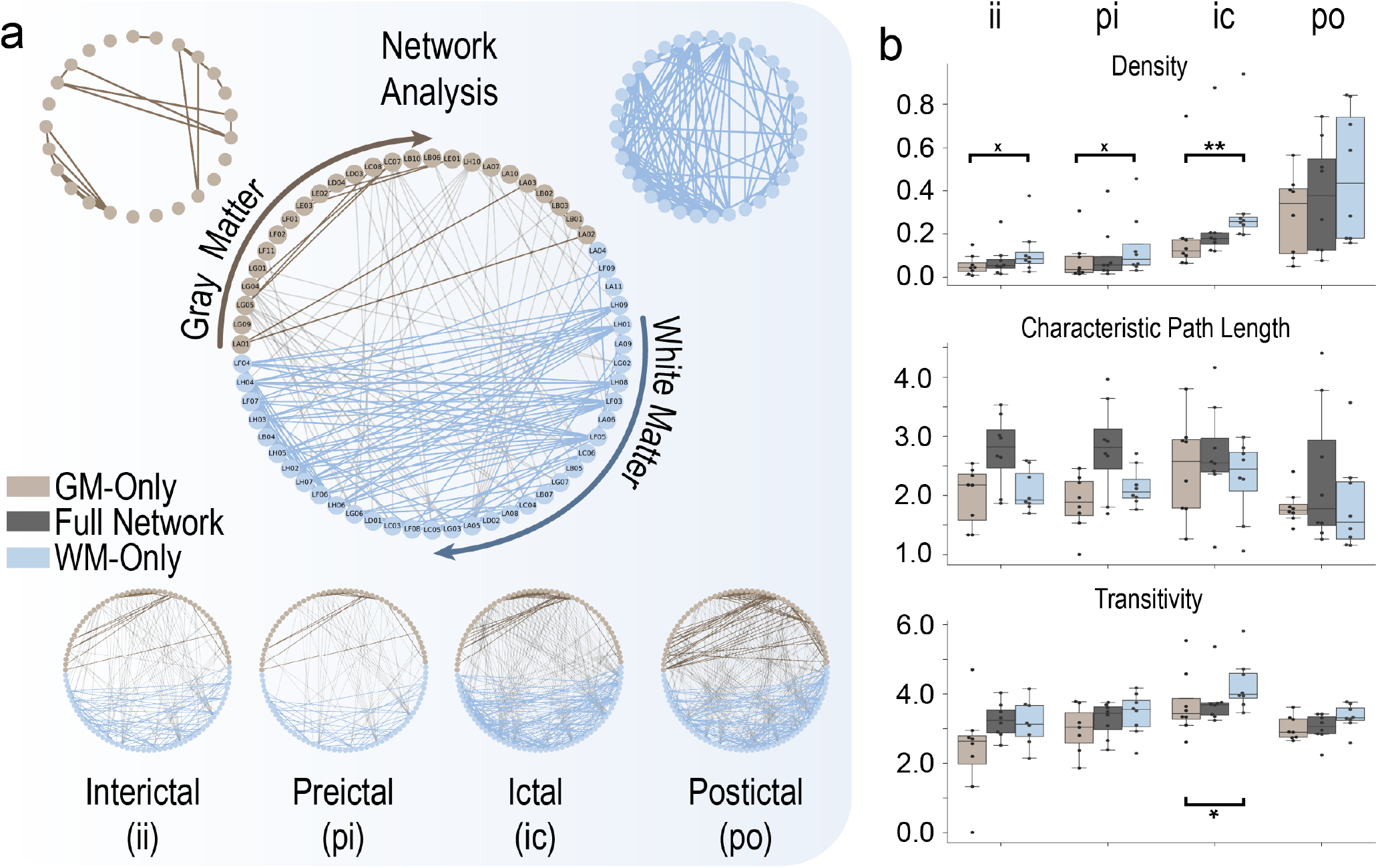
White matter (WM) tissue sub-networks have different network properties than gray matter (GM). **a**, Network visualization of an example patient during the ictal state. WM contacts are ordered clockwise by increasing WM distance from GM contacts. Another example patient during all four states is shown (bottom), illustrating the increase in functional connectivity during the ictal state. **b**, Network measures of density, characteristic path length, and transitivity during each state are plotted. Density and transitivity are higher in WM than GM tissue sub-networks (*p*< 0.05); however, the characteristic path length is not different. WM networks are denser and more clustered. Density and transitivity is higher in WM than in GM sub-networks for the interictal and preictal state. (*trending, p*< 0.1 indicated by an *x*).

### White matter recordings capture the communication between gray matter regions during seizures

The previous analyses above illustrate the altered signaling and seizure dynamics of GM and WM recordings: WM signals have increased FC during seizures. We next sought to explain what drives this activity. Does this increased WM FC result from the activity of other GM tissue (for example, from the SOZ), or is the increase in FC intrinsic to WM tissue itself during the ictal state? We applied a structure-function analysis by assessing the statistical correlation between structural and functional connections to determine if the increased FC in WM is explained by the structural connections from GM. If so, we and others hypothesize that WM signals reflect the communication between GM regions^7,8^. In patients with high angular diffusion imaging (HARDI, *N* = 16), we measured the structural connectivity between contacts (Fig. 4a). We also show the distribution of functional connectivity values in Fig. 4b. We calculated the structure-function correlation (SFC) as a correlation coefficient between the upper triangle of the structural matrix and the upper triangle of the functional matrix, separately for the four tissue networks (Fig. 4c). We show that the SFC increases for the full, GM-only, and GM-WM networks (*p* < 0.05), but not for the WM-only sub-network (Fig. 4d, top). We bootstrapped our data 10,000 times (Fig. 4d, bottom) and calculated the mean SFC for each simulation. We also calculated the change in SFC (Δ*SFC*) from preictal to ictal states (Δ*SFC* = *SFC*_*ic*_ − *SFC*_*pi*_) and plot the distributions of Δ*SFC* over 10,000 bootstrap simulations (Fig. 4e). The SFC increases the most from preictal to ictal states for GM-WM sub-networks and the least for WM-only sub-networks (Δ*SFC* means with 95% confidence intervals: Full, 0.0537 [0.0529-0.0544]; GM-only, 0.0525 [0.0516-0.0533]; GM-WM, 0.0771 [0.0758-0.0783]; WM-only, 0.0312 [0.0299-0.0324]). These results support the hypothesis that the change in WM at seizure onset is driven by the structural connections from GM regions (e.g., the SOZ or other regions involved in seizure activity, Fig. 4f).

**Fig. 4.**
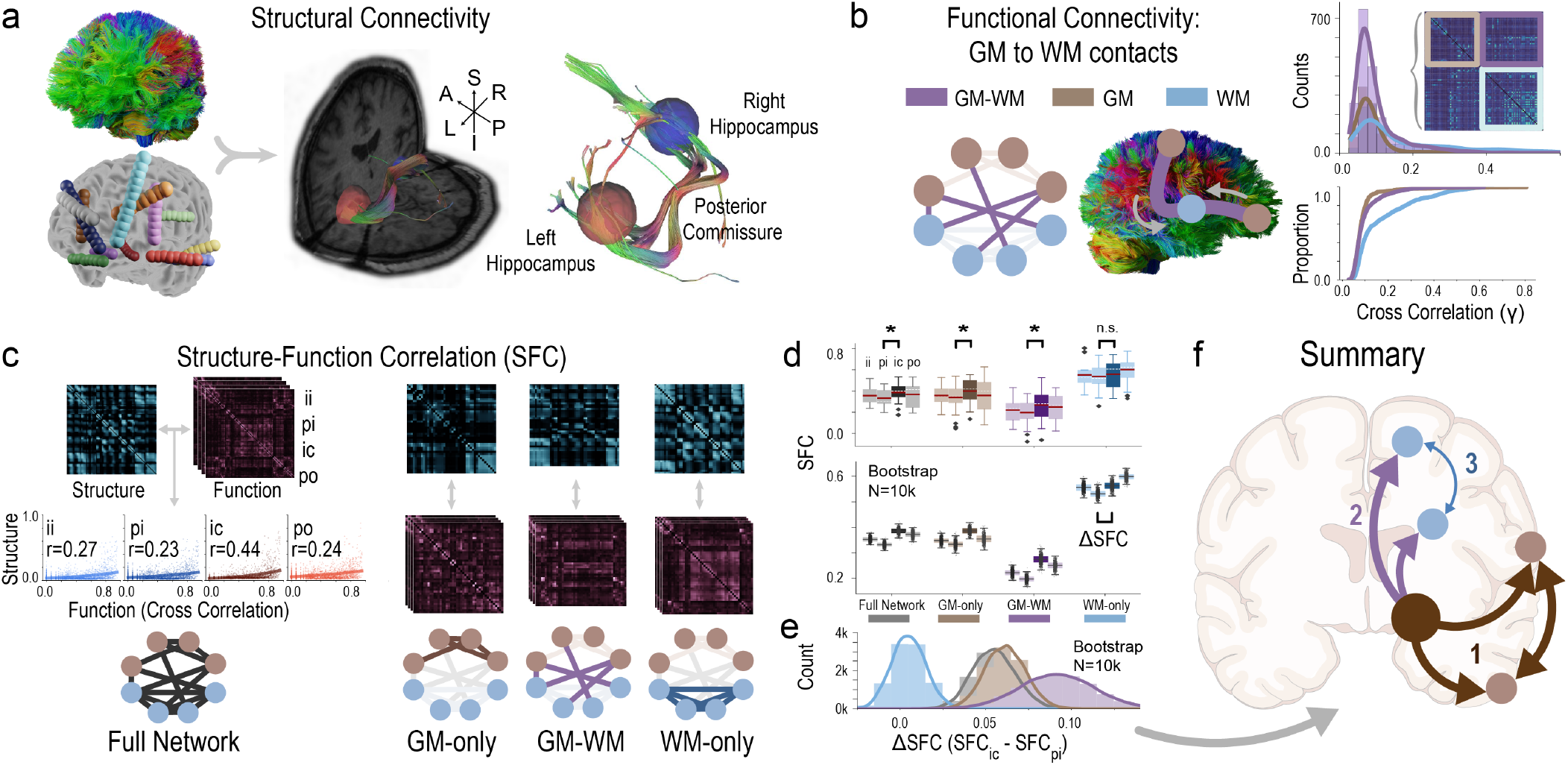
White matter (WM) recordings capture the communication between gray matter (GM) regions. **a**, WM tracts are visualized between the left and right hippocampi for an example patient. **b**, Histogram distributions and ECDFs are shown for the different tissue sub-networks similar to Fig. 2c. **c**, Structure-function correlation (SFC) is computed for each sub-network. *r* = Spearman Rank correlation. **d**, SFC for each state and sub-network is shown (top). Preictal and ictal SFCs are compared. Bootstrapped means (*N* = 10, 000) of SFC for each state and each sub-network is shown (bottom). **e**, The distribution of bootstrapped means (*N* = 10, 000) of Δ*SFC* (*SFC*_*ictal*_ − *SFC*_*preictal*_) is plotted showing 6-*SFC* is highest for GM-WM connections and lowest for WM-only sub-networks. **f**, Summary of findings: (1) GM functional activity changes during seizures are correlated to other GM regions (and are documented through other studies^15,16^). (2) WM functional activity changes during seizures are correlated to the structural connectivity from GM regions. (3) WM functional activity changes during seizures have the lowest correlation to the structural connectivity from other WM regions (see confidence intervals in the main text). This observation supports the notion that activity in WM is driven by activity in GM and may be explained by the structural connections between regions. **SFC**, Structure-Function Correlation.

### Surgical epilepsy patients with poor outcomes have higher ictal SEEG white matter connectivity than patients with good outcomes

A total of 17 patients had outcome scores ≥ 2 years after epilepsy surgery to remove the SOZ (good outcome *N* = 10; poor outcome *N* = 7). Chi-squared test with Yates’ correction shows non-statistically significant differences in good versus poor outcome epilepsy patients with respect to epilepsy type (MTLE versus Non-MTLE, Chi squared = 0.084 with 1 degrees of freedom; two-tailed p > 0.05, Table S1). Similarly, Chi-squared test with Yates’ correction shows non-statistically significant differences in good versus poor outcome epilepsy patients with respect to lesional status (Chi squared = 0.093 with 1 degrees of freedom; two-tailed p > 0.05).

We show that poor outcome patients have a stronger in-crease in WM FC during the ictal state (see overview in Fig. 5a). During the ictal state, the full, GM-WM, and WM-only sub-networks have higher connectivity (*p*< 0.05) in poor outcome patients than in good outcome patients (Fig. 5b). This increase is not apparent in the GM-only sub-network, which is the sole network available for patients that only have GM implanted or for analyses that only consider GM contacts (*p* > 0.05). We repeated the analysis in Fig. 2h, separating good and poor outcome patients (Fig. 5c). We show that the differences between GM and WM sub-networks during the ictal period become more pronounced for poor outcome patients than for good outcome patients using re-sampling hypothesis testing (*p* < 0.05, Fig. 5d, top) and bootstrapping 10,000 times the mean Δ*r*_*ictal*_ of good and poor outcome patients (Fig. 5d, bottom).

**Fig. 5.**
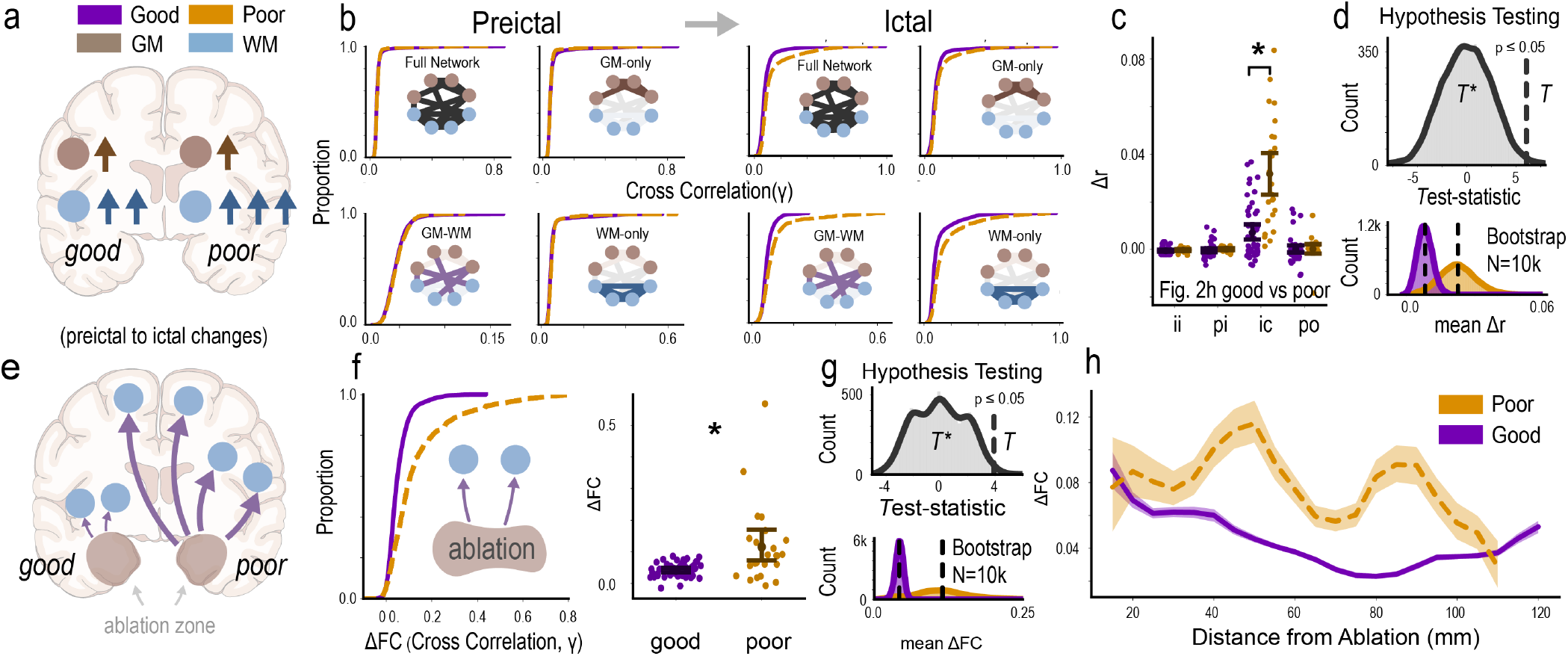
Good outcome epilepsy patients have lower white matter (WM) functional connectivity than poor outcome patients. **a**, Diagram summarizing results for the top row: WM and GM FC increase during seizures for both good and poor outcome patients. WM FC increases more than GM FC (see Fig. 2i), and this increase is even larger for poor outcome patients. **b**, Empirical cumulative density functions (ECDF) are shown for the sub-networks in poor and good outcome patients. FC is higher during the ictal state for poor outcome patients in the full, GM-WM, and WM-only tissue sub-networks. However, FC is not significantly different for the GM-only tissue sub-network. **c**, Differences in GM and WM FC values (Δ*r*) during the ictal period are larger for poor outcome patients than for good outcome patients, consistent with Fig. 2h. Error bars represent 95% confidence intervals. **d** Re-sampling hypothesis testing (top) confirming the analysis of Fig. 5c (see the re-sampling hypothesis testing subsection of the Methods). Dashed vertical line: *T*. The distribution of bootstrapped means of the ictal Δ*r* is also plotted comparing good and poor outcome patients (bottom). **e**, Diagram summarizing results for the bottom row: ablated regions in good outcome patients have lower changes in FC to distant WM contacts than poor outcome patients during seizures. **f**, ECDF of the FC values between the contacts in the ablation zone and all WM contacts (left). The median FC values for each re-sampled patient is compared (right). **g**, Same as in panel **d** but now comparing the changes in FC between ablated and WM contacts. **h**, The FC values between the contacts in the ablation zone and WM contacts as a function of distance. We show that the FC of ablated contacts to distant WM contacts increases more during seizures for poor than good outcome patients ***T***^***\****^, test statistic of permuted samples; ***T***, test statistic of original data.

### Ablated regions in good outcome patients have lower ictal connectivity to white matter signals than poor outcome patients

In both good and poor outcome patients, the ablation area was targeted because its removal would give the best chance of seizure freedom despite the surgery’s inherent risk. Surgery was performed after a comprehensive clinical assessment of data acquired from multiple sources including various MRI sequences, scalp EEG, iEEG, patient and family histories, and neuropsychological evaluation. We showed in Fig. 5 (top row) that differences in FC of good and poor outcomes are mainly driven by connections to WM, but not apparent in GM-only regions. We tested if the FC between ablated regions and WM contacts could differentiate between good and poor outcome patients (Fig. 5e). We show that FC between the ablated regions and WM contacts is higher for poor outcome patients than good outcome patients (*p* < 0.05, Fig. 5f-g). This finding may indicate that the suspected SOZ in poor outcome patients (which would eventually be ablated) is not a good target given that its activity has a high correlation with the activity of WM contacts.

### White matter functional connectivity changes during seizures are elevated far from the ablation zone in poor outcome patients

We next sought to determine whether the correlations remain elevated in WM contacts distant from the ablation site in poor outcome patients but not in good outcome patients (Fig. 5h). After bootstrap re-sampling of patients and seizures, we show that WM FC to the contacts contained in the *eventual* ablation zone remains elevated in poor outcome patients. This increase is most apparent for WM contacts 40–60 mm (approximately a half hemisphere away) and 80–100 mm (approximately on the contralateral hemisphere) from the ablation zone. Future analyses should further contextualize the location of these WM contacts to canonical WM pathways.

## Discussion

We demonstrated that white matter intracranial EEG (iEEG) recordings reveal underlying seizure dynamics that are not captured in gray matter recordings alone. We compared gray matter and white matter recordings from stereoelectroencephalography (SEEG) implantations of 29 medically refractory epilepsy patients and found differences in signal and network properties of white matter contacts (Figs. 1-3). We showed that white matter contacts capture neural communication along the structural pathways connecting brain regions in 16 of the SEEG patients who acquired high angular resolution diffusion imaging (HARDI, Fig. 4). These white matter recordings also have different seizure dynamics in good and poor outcome epilepsy patients at distant sites from the ablated regions (Fig. 5). Our study provides evidence that the analysis of white matter functional signals may prove worthwhile when researching and developing new diagnostic and treatment strategies for medically refractory epilepsy patients. More broadly, white matter functional signals may also reveal important clues to brain function and pathophysiology of other neurological disorders. To facilitate reproducibility and further investigation of these SEEG patients, we have made the iEEG data, metadata, and de-identified processed structural tractography neuroimaging data compliant with Brain Imaging Data Structure (BIDS) and publicly available (see Methods and ‘Data availability and reproducibility’).

### Biological processes revealed in white matter

Historically, neuroscientists have “exhibited little interest in white matter”^20^ because it was thought to be an inert, passive tissue, despite comprising approximately 50% of the brain’s volume^14,21^. Instead, investigators have focused on gray matter functional activity because neural computation primarily takes place in gray matter, and gray matter signals can be orders of magnitude more powerful than white matter^7,8,13^. Only recently has white matter been appreciated to support many critical brain functions^22^, modulate the distribution of action potentials^23^, and act as a relay of neural communication between different brain regions^24^. Notably, mounting evidence confirms that neuronal cell bodies exist in deeper white matter tissue (>2–3mm from the cortical layers)^25^; neurotransmitter vesicles are released directly in white matter, affecting neural communication^26^; and white matter blood-oxygenation level dependent (BOLD) signals are detectable across a range of different tasks^27^. All of these findings reflect white matter is indeed a dynamic, active tissue and its recordings may reveal important clues to brain function and pathophysiology^28^.

These findings raise several questions. What are the functions of white matter neurons in epilepsy? What are the consequences of altered neural communication resulting from the release of neurotransmitters along the axons of white matter? Can white matter activity recorded by fMRI, iEEG, and other technologies reveal clues to epilepsy pathophysiology and, more generally, neurophysiology?

Here, we have provided examples of white matter’s utility for neuroscience inquiry through a multi-pronged approach (from signals to networks) across many brain states (from interictal to postictal) and with multimodal data (from structure to function) in different outcomes of epilepsy patients (from good to poor). We provide ample evidence highlighting the importance of white matter signals, which could be further investigated in future studies to reveal novel insights into neurobiology, disease processes, and diagnoses.

### Intrinsic or extrinsic signals of white matter? Capturing information transmission between brain regions

White matter is considered a structural tissue that transmits neural information from one gray matter region to another, yet it is unknown what the *functional* signals captured in white matter (namely, the iEEG signals from WM contacts in our study) represent. If the signals were merely local field potentials (LFPs) generated by local neurons^25^ that were affected by the release of neurotransmitter vesicles in white matter^26^, there would need to be a sufficient number of geometrically aligned neurons to produce these fields. Evidence of detectable white matter BOLD signals in fMRI may support this “intrinsic” activity of white matter^27^. However, our results support that the activity captured from within the WM (at least, electrical – not metabolic) originates extrinsically, and probably not solely due to volume conduction of nearby gray matter tissue (Fig. 4). We show across multiple frequency bands and montaging schemes that white matter signals are related to their direct structural connections from other gray matter tissue (Fig. 4e), supporting that these signals may capture the transmission of information between brain regions (Fig. 6). In other words, gray matter signals drive white matter signals. Thus, these signals may be considered more extrinsically driven. The “intrinsic” tissue metabolism detected in these regions may be extrinsically related at some level, too.

**Fig. 6.**
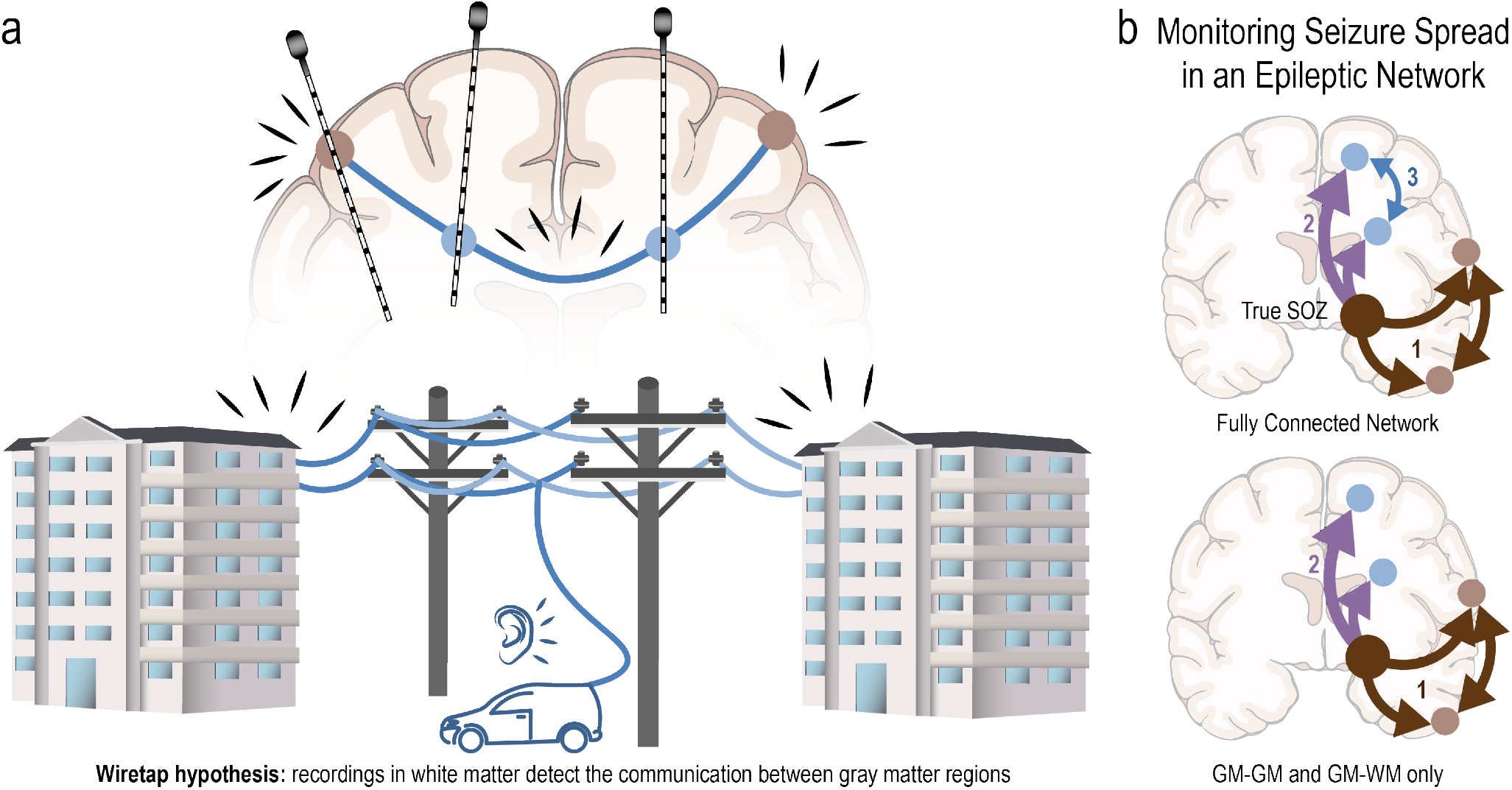
White Matter Signals Reflect Information Transmission Between Brain Regions During Seizures. **a**, Showing that white matter recordings capture the communications between gray matter regions has implications in localizing the seizure onset zone in epilepsy patients. For example, an apt analogy could be made between white matter recordings and monitoring (or “tapping”) the activity on a phone wire between two buildings (the “wiretap hypothesis”). Information about the phone conversation is acquired without ever having to monitor the locations where the conversation is physically happening. It is especially challenging in epilepsy to precisely locate the seizure onset zone in the face of imperfect implantation schemes in which we cannot sample all brain regions. Here, we allude to the fact that white matter recordings may provide information about the seizure onset zone without ever having to implant the seizure onset zone itself, because white matter recordings monitor the activity between gray matter regions. All we need to know is where the connections are (or in our phone analogy, which buildings the phone wires go to). This information can be acquired more easily though non-invasive neuroimaging. In practical terms, if we were able to detect specific ictal signals within WM contacts, finding which GM structures are connected through those specific WM fibers could reveal information about the SOZ. **b**, In analyzing the spread of seizure activity from contact to contact, it would be helpful to know the gray matter regions to which the white matter contacts are structurally connected (connections 1 and 2), but not necessarily which to which other white matter contacts they are connected (connection 3). To predict seizure spread, in other words, white matter activity should be predicted from gray matter activity, but not from other white matter recordings.

Showing that white matter recordings can capture communications between gray matter regions has profound implications for localizing seizure onset zones in epilepsy patients. For example, an apt analogy could be made between white matter recordings and monitoring (or “tapping”) the activity on a phone wire between two places (Fig. 6a). Information about the phone conversation is acquired without ever having to monitor the locations where the conversation is physically originating or being received. It is especially difficult in epilepsy to precisely locate the seizure onset zone in the face of SEEG implantation schemes since sampling of all possible brain regions is not feasible. Here, we propose that white matter recordings may provide information about the seizure onset zone even if the seizure onset zone itself were not targeted in the implantation. Because white matter recordings reflect the activity between the gray matter regions that they link, one could infer which two structures are connected (or in our phone analogy, to which buildings the phone wires go) by identifying the ends of the white matter fiber tract showing aberrant activity. Indeed, this information can be acquired more easily though non-invasive neuroimaging, supplementing the information gathered through iEEG. Furthermore, in analyzing the spread of seizure activity from contact to contact, it would be helpful to know which gray matter regions the white matter contacts are structurally connected to, but not necessarily to which other white matter regions they are structurally connected to (Fig. 6b). To predict seizure spread, in other words, white matter activity should be predicted from gray matter activity, but not from other white matter recordings.

### The Distributed Epileptic Network Hypothesis

We provided evidence that white matter recordings reveal differences in epilepsy pathophysiology of good and poor outcome epilepsy patients (Fig. 5). Poor outcome patients have higher FC between the *suspected* (and eventually ablated or resected) SOZ and more distant white matter recordings than good outcome patients (Fig. 5h). We attribute these differences to a broadly distributed (i.e., non-focal) seizure etiology of poor outcome patients. The areas ablated in poor outcome patients were chosen because of supporting clinical evidence that the primary target for ablation was a well-localized, focal SOZ; otherwise, patients would not have been deemed candidates for surgery. Yet, patients continue having seizures after ablation. The inefficacy of the surgery could be due to at least two factors: (1) the SOZ is still a focal region, but was missed in the ablation/resection, or (2) the SOZ appears focal to clinicians, but the underlying pathophysiology in some patients is due to a more widely distributed interaction of brain regions and interconnecting white matter^29^. Note that in the first case, patients with a single epileptic focus may still have widespread structural, functional, or behavioral (i.e., clinical) changes and/or deficits resulting from their seizures, perhaps due to aberrant plasticity over time^30^. However, in the second case, the etiology (rather than the *sequelae*) stems from an “epileptogenic”^31^ or “epileptic”^5^ network of interacting brain regions, allowing seizures to occur.

The traditional model of the epileptogenic focus is too simplistic to capture the spatiotemporal organization of seizures^32^. Thus, many investigators have utilized complex system approaches instead, showing that epilepsy may be a disease of networks with structural (e.g., DTI), functional (e.g., fMRI), and effective (e.g., CCEPs) data^15,33–39^. Here, we provided evidence for the epileptic network captured in the white matter of poor outcome patients. We build on the concept of the epileptic network to explain *why* some patients have poor outcomes. We hypothesize that, in these patients, the true SOZ is not a single focal anatomical location, but rather a *distributed epileptic network* where many brain regions interact, causing patients’ neural dynamics to bifurcate^40^ or spontaneously devolve into seizure states. We take the opportunity to name the hypothesis we proposed in the previous sentence the “**Distributed Epileptic Network Hypothesis**” (Fig. 7) to provide concrete language for discussing the pathophysiology of some epilepsy patients, in hopes of spurring new areas of research and therapeutic treatment options for these patients. In these poor outcome patients, important nodes of the epileptic network were eliminated during surgery; thus, they may be considered Engel III with “marked seizure reduction” and reduced disease burden. However, seizures eventually reoccur because the epileptic network was not entirely hindered. Instead, only select important nodes were eliminated. The network then reacts or it reforms^41^ to produce seizures, albeit not as strong or as frequent before surgery (Engel II/III). Here in our paper, we show evidence for a more broadly distributed epileptic network in poor outcome patients that is captured in white matter recordings.

**Fig. 7.**
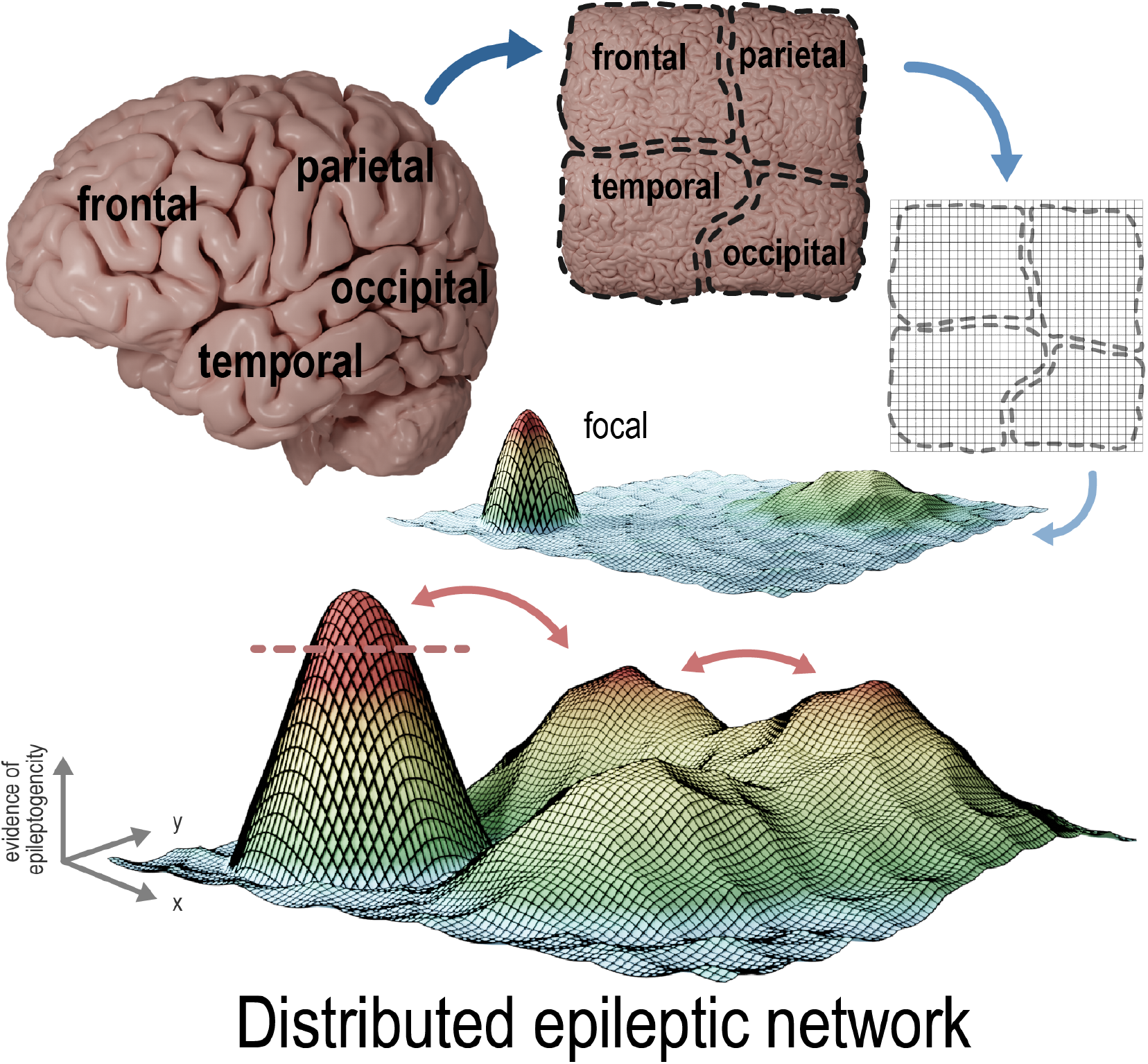
The Distributed Epileptic Network Hypothesis. A distributed epileptic network can arise when multiple brain regions *interact* together to produce seizures, rather than one singular focus causing seizures. This interaction between brain regions may explain why, in some patients, seizure freedom is not achieved even in the setting of *focally-appearing* epilepsy. Here, the red dotted line represents a cutoff where, given evidence from pre-surgical diagnostic testing, a single epileptogenic region presents itself as seizure onset to a team of physicians. But in reality, elimination of this region may not yield seizure freedom, or seizure freedom only for a short period of time. Distributed regions of the brain interact, adapt, and reform to produce seizures, even if an important node has been eliminated. The landscape of epileptogenicity (the illustrated mesh) can change over time. In this manuscript, we show that evidence of a distributed epileptic network may appear in the recordings of white matter. Electrodes inserted into the white matter tracts across distributed regions of the brain have larger increases in correlations during seizures to the ablation/resection site of poor outcome patients than good outcome patients (Fig. 5f).

### Limitations

Our study is not without limitations. They include: (1) The choice of montaging and frequency bands. We attempted to extract the intrinsic white matter signals or other signals coming from distant brain areas, and not from volume conduction of nearby gray matter. Regardless, volume conduction from other confounding extrinsic sources cannot be eliminated completely and may increase FC between certain brain regions, thereby affecting our findings. (2) A small sample size of poor outcome patients and definitions of outcomes. We aimed to mitigate the biases of patients, different implantation schemes, and different tissue sampling techniques through bootstrapping, another approach using re-sampling hypothesis testing, and more stringent exclusion criteria of outcome ≤2 years. (3) Seizure temporal dynamics. We computed FC over entire segments of interictal, preictal, ictal, and postictal states. Seizure dynamics are not the same between early and late stages of seizure evolution. FC differences between white matter/gray matter and good/poor outcome patients could lead to a different interpretation when involving more complex time-dynamical analyses. (4) White matter location. We did not include an analysis of the anatomical structures of white matter contacts or test if specific structures are mainly driving white matter FC (possibly supported by Fig. 5h). For example, we did not analyze the corpus callosum or corona radiata fibers that are typically captured in SEEG implantations along the trajectory to deeper gray matter targets. Note that the limited research of WM activity in fMRI has focused on the corpus callosum^42^. (5) Sampling technology. Other implantable devices such as microelectrode-arrays (MEAs) or Utah arrays^43^, which can record at higher densities, altered to target white matter may provide more robust findings about white matter signals. In contrast, SEEG records LFPs at the meso-scale.

### Future Directions

White matter functional recordings acquired through many technologies (including iEEG, fMRI, MEG, PET, ASL, and others) may provide a wealth of *currently untapped* knowledge about neurobiology and disease. Our work has ramifications across the field of neuroscience and highlights the possibility of new areas to explore: stimulation of white matter to modulate brain function; biomarkers for monitoring white matter functional signals in the progression, and evolution of, other neurological and psychiatric diseases such as multiple sclerosis, dementia, and schizophrenia; the utility of targeting white matter structures directly, rather than coincidentally acquiring signals in targeting gray matter structures; direct ablation or surgical intervention of white matter areas to improve outcomes, rather than focusing on gray matter structures only; drug therapies that focus on the modulation of abnormal white matter activity, rather than using gray matter activity benchmarks. More research is needed to assess the viability of all these options. The focus of treatment strategies can be on white matter, too.

## Supporting information

Video Abstract

## Materials and Methods

### Human Dataset

Twenty-nine drug-resistant epilepsy patients underwent stereoelectroencephalography (SEEG) implantation for clinical purposes at the Hospital of the University of Pennsylvania. Of these patients, 88 seizures were recorded, with distributions of seizures per patient and seizure lengths shown in Fig. S5. Sixteen of the SEEG patients had MRI data collected for SFC analyses. Inclusion criteria consisted of all individuals who agreed to participate in our research scanning protocol, and allowed their de-identified intracranial EEG (iEEG) data to be publicly available for research purposes on the International Epilepsy Electrophysiology Portal (https://www.ieeg.org)^44,45^. Seizure evaluation was determined via comprehensive clinical assessment, which included multimodal imaging, scalp and intracranial video-EEG monitoring, and neuropsychological testing. This study was approved by the Institutional Review Board of the University of Pennsylvania, and all subjects provided written informed consent prior to participating. See Table S1 for subject demographics. Sample sizes were determined by attempting to gather as many patients as possible with validated outcome scores ≥ 2 years, annotated seizure markings, and high angular resolution diffusion imaging (HARDI). No statistical methods were used to predetermine sample sizes, but our sample sizes are larger than those reported in previous publications using both iEEG and diffusion imaging data^15,16,46^

### Intracranial EEG Acquisition

Stereotactic depth electrodes were implanted in patients based on clinical necessity. Continuous SEEG signals were obtained for the duration of each patient’s stay in the epilepsy monitoring unit. Intracranial data was recorded at 256, 512, or 1024 Hz for each patient. Seizure onset times were defined by the unequivocal electrographic onset (UEO)^47^. All annotations were verified by neurologists and consistent with detailed clinical documentation. The spacing between SEEG contacts is 5 mm and the contacts are 2.41 mm in size.

### Electrode Localization

See Fig. S1 for an overview of electrode localization. In-house software^48^ was used to assist in localizing electrodes after registration of pre-implant and post-implant images (T1w and CT images). All electrode coordinates and labels were saved and matched with the electrode names on IEEG.org. All electrode localizations were verified by a board-certified neuroradiologist (J.S.).

### Tissue segmentation

See Fig. S1 for an overview of tissue segmentation. Freesurfer recon-all tissue segmentation output was used. The pre-implant T1w image specified in the imaging protocol below was used for tissue segmentation. If no T1w image from this protocol was acquired, the T1w image from the pre-implant clinical scan was used. The tissue segmentation (the “aseg” file output from Freesurfer) file which includes cortical GM, subcortical GM, WM, and other anatomical labels was converted to 4 classes: GM, WM, CSF, and outside. All cortical and subcortical GM structures were combined to a single GM class. WM structures were all combined to a single WM class. Voxels contained within the Freesurfer brain extraction file, but outside GM or WM structures, were labeled as CSF. All other voxels not in WM, GM, or inside the brain extraction file were labeled as outside the cranium.

### Contact tissue localization

The post-implant T1w image used for electrode localization underwent linear registration to the preimplant T1w image used for tissue segmentation above. Linear registration was performed using *fnirt* in the FSL neuroimaging software. Contacts were classified to the tissue classes defined in the section below. Only contacts where the center coordinate fell into GM or WM labeled voxels were used. CSF labeled contacts, which generally included those between the skull and brain, and all other contacts were excluded from analysis. Note that for tissue localization purposes, implanted contacts were rarely localized to non-mesial temporal subcortical structures (i.e. the thalamus, putamen, etc.). These structures are rarely targeted in epilepsy patients. Thus accurate tissue segmentation of these subcortical structures were not vital for our study.

### GM and WM definitions by distance or depth

Contacts localized to WM voxels may not necessarily record exclusively from WM, especially when neighboring voxels are GM cortical structures. This was addressed in Mercier et al. 2017 using a measure called proximal tissue density (PTD)^7^. This “fuzzy” approach is used to (1) mitigate volume conduction (2) mitigate the degree of uncertainty in the location of the contact (can be due to small errors in registration). Here, we follow a similar fuzzy approach to define GM and WM contacts. See Fig. S2 for an overview.

Definitions:

1. **Distance**: Distance to the nearest GM voxel. Typical range 0-10mm. Contacts that were 0mm away (i.e. the contact centroid coordinate was in a GM voxel) were labeled as GM. Analyses that relied on a definition of distance used the following criteria:
  a. GM: = 0 mm
  b. WM: ≥ 2 mm
  c. contacts between 0 and 2mm were excluded from analysis comparing GM and WM. This follows the Mercier et al. 2017^7^ approach where PTD the extreme values of -1 and 1 were used.
2. **Depth**: Percent of surrounding voxels that are WM. Typical range: 0-1 (0-100%). A higher WM contact depth equates to a contact deeply embedded into WM tissue with little surrounding GM tissue. This convention is similar to the proximal tissue density (PTD) used in Mercier et al. 2017^7^. A sphere with a diameter of 1cm around the centroid coordinate of the contact was used to calculate the percent WM contained within that sphere. Analyses that relied on a definition of depth used the following criteria:
  a. GM: ≤ 0.5 (or 50%).
  b. WM: ≥ 0.9 (or 90%)
  c. contacts between 0.5 and 0.9 were excluded from analysis comparing GM and WM

The above definitions, along with bipolar montaging (below) help ensure the recorded signals are a good representation of activity of that tissue and contact location. We chose a definition of WM depth for figure analyses in the main text because it is similar to PTD, unless specified WM distance was used (i.e. power analysis vs distance and SNR in Fig. 1c, and controlling for WM distance in Fig. 2f). Please see the main text for a discussion on why two definitions were used.

Briefly, alternative definitions provide different perspectives when each respective tissue definition may have its own limitations (Fig. S2). For example, definition 1 (distance) may incorrectly say a WM contact is close to GM because a single GM voxel was mislabeled due to imperfections of the Freesurfer tissue segmentation or due to small errors in registration. Regardless, previous studies^8^ and our results in “Controlling for distances between contacts” show similar findings using alternate definitions. For subsequent analyses, we used definition 2 (depth) because it is a similar and complimentary definition to PTD used in Mercier et al. 2017^7^, yet still provides a different perspective in this nascent field for understanding WM iEEG activity where a standard WM iEEG definition is not set. The boundaries for classifying GM and WM for each definition (e.g. ≥ 2 mm) was slightly arbitrary and set before conducting the study: two millimeters for definition 1 was chosen because that is approximately where “deep” white matter is defined in the histological field^25^. For definition 2, 50% for GM was chosen because that seemed appropriate where the minority of surrounding voxels are WM. Similarly, 90% for WM was chosen because that seemed appropriate where extreme values of PTD were used in prior studies. Future studies should explore these exact parameters, however, it is out of the scope of this study given that our work and other groups have already shown that the nuances of tissue definitions are not necessarily driving the biological findings. After peer review, we added another analysis based on the amount of gray matter surrounding a contact (rather than based on white matter above). Results are similar and presented in Fig. S10.

### Contact localization to ablation zone

Contacts contained within the eventual ablation zone were manually identified. Post-surgical T1w images were acquired and manually segmented using ITK-SNAP for the areas ablated. The post-surgical T1w image was linearly registered to pre-implant image. Contact coordinates were found in pre-implant space (electrode localization pipeline above). Contacts that fell into the ablation segmentation mask were labeled as “ablated contacts.”

### Montaging

While epileptologist routinely use bipolar montage in EEG interpretation, the research community does not. A few studies^7,8^ noted differences in EEG features like power, phase-amplitude coupling, and functional connectivity measurements between the different montages. Makin et al. 2020^49^ emphasize that if the electrodes are common average referenced, it can “smear the effects of some electrodes across the grid”. Bipolar referencing was noted to improve overall performance using their deep learning approach. Here we use bipolar referencing to mitigate the effects of volume conduction, not in line with previous studies using common average referencing to create functional connectivity matrices to study epilepsy. To perform bipolar referencing, we started with the contact at the most distal end of the electrode (e.g. LA01) and subtracted the adjacent contact on that electrode (e.g. LA01 = LA01 - LA02). We excluded the last contact on the electrode (there is nothing to subtract), or any bipolar pairs that included a contact excluded through any methods mentioned throughout this methods section (e.g. an artifact contact or a contact not in GM or WM). A critical note is that bipolar montaging, along with the GM and WM contact definitions (above) help ensure the recorded signals are a good representation of activity of that tissue and contact location. Each montage has their own respective strength and weaknesses, which is why we opted to show results using common average reference (CAR, Fig. S8). We feel that providing both perceptive with repeatable results strengthens our conclusions.

### Power Spectral Density Calculation

Following removal of artifact-ridden electrodes, SEEG signals were bipolar referenced. Signals were notch-filtered at 60 Hz to remove power line noise and low-pass and high-pass filtered at 127 Hz and 1Hz to account for noise and drift. Power spectral density (PSD) was calculated using the Scipy Python package version 1.5^50^, and the function scipy.signal.welch (default parameters, with FFT epoch length equal to 1s). For the power vs distance analysis, power was taken over the entire state (interitcal, preictal, ictal, postictal). For the SNR analysis, power was taken over 1s intervals. Power was log10 transformed on a *V*^2^*/Hz* scale.

### Signal-to-Noise Ratio Calculation

Here we define signal-to-noise ratio (SNR) as the ratio of the ictal power (‘signal’) to interictal power (‘noise’). This definition follows the framework in which epileptologists compare background interictal or preictal activity with ictal activity using EEG features such as amplitude, frequency, and timing. Comparison of any epileptiform signals to this background noise aids in seizure diagnosis and localization.

Over each state, we normalized all times to a length of 100. For example, all interictal, preictal, and posictal segments were 180 seconds (see functional connectivity network generation below). The SNR vectors (180 one-second intervals long) were normalized to a length of 100 for each patient. For ictal segments of variable length of each patient, vectors were normalized to a length of 100. Thus 50% on the x-axis of Fig. 1c represents half-way through each segment (i.e. 90s into the interictal segment, or 50s into a 100s long seizure). Patients’ SNR was averaged and plotted. For the GM vs WM SNR comparison, the boxplot of all patients’ SNR was plotted at 50% into each state. We then performed a Mann-Whitney U rank test to compare the SNR of GM (distance from GM = 0) and WM ≥6mm from GM at the 50% time interval.

### Functional Connectivity Network Generation

Functional connectivity networks were generated from four states: interictal (ii), preictal (pi), ictal (ic), and postictal (po). (1) The interictal state consisted of the time approximately 6 hours before the ictal state. (2) The preictal state consisted of the time immediately before the ictal state. (3) The ictal state consisted of the time between the seizure unequivocal onset and seizure termination. (4) The postictal state consisted of the time immediately after the ictal state. Interictal, preictal, and postictal periods were 180 seconds in duration. To generate functional networks defined by cross correlations between SEEG signals, we detail our methods below, in line with previous publications that calculate functional connectivity on iEEG data^15,17^. SEEG signals were preprocessed using methods outlined in the PSD section above (i.e. artifact removal, bipolar referencing, and filtering). Signals were then pre-whitened using a first-order autoregressive model to account for slow dynamics. Functional networks were then generated by applying a normalized cross correlation function *ρ* between the signals of each pair of electrodes within each of the four states, using the formula:

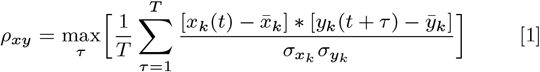

where x and y are signals from two electrodes, k is the state, t is one of the T samples during the time window, and *τ* is the time lag between signals, with a maximum lag of 0.5 s. The absolute value of the cross correlation values were taken. For more details, please see prior publications^15,17^.

### Frequency band definition

Frequency bands were defined using typical ranges reviewed in Newson and Thiagarajan 2019^51^, rounded to the nearest integer:

1. Broadband: 1-128Hz
2. *δ* (delta) band: 1-4Hz
3. *θ* (theta) band: 4-8Hz
4. *α* (alpha) band: 8-13Hz
5. *β* (beta) band: 13-30Hz
6. *γ*(gamma) band: ≥ 30 Hz
  a. low-*γ*30-40Hz
  b. mid-*γ*70-100Hz
  c. high-*γ*100-128Hz

The frequency band with the widest range reported in Newson and Thiagarajan 2019^51^ after reviewing 184 studies was the *γ*-band. A typical *γ*-band was 30-40 Hz. We subdivided this band further^15,17^. Low-*γ*equates to the typical *γ*-band. Mid-*γ*skips the 60Hz power line noise (or any potential imperfections in its elimination) and starts at a lower bound of 70Hz. High-*γ*starts at 100Hz and the upper limit was chosen as 128Hz because that was half the sampling frequency of the patient with the lowest sampling frequency. We chose to analyze the high-*γ*frequency band because of minimal volume conduction, and well-developed literature studying high-*γ*in epilepsy (however, it is less consistently defined across the literature, but typically >100Hz). Unfortunately, we could not analyze higher frequency bands due to non-uniform sampling frequencies across all patients.

An important note is we show consistent results over many different frequency bands. Differences in frequency band definitions across studies have enormous consequences for interpretation, which is why we opted for a review that performed a meta-analysis on a large number of studies (N = 184 studies) by Newson and Thiagarajan, 2019^51^. Even if the study is from outside of epilepsy, the work they did still provides utility to the neuroscience, neurology, and psychiatry fields at large. An important concern with any study is that differences found may disappear when moving the band windows slightly. However, we show that results are repeatable at many frequency bands, not only with our “high” gamma reported in the main text. A wider frequency band for “high” gamma of approximately 70 — 150Hz may be more widely used, as with Kucyi et al. using 70 – 170 Hz^52^, Grossman et al. using 45-154 Hz^53^, and Mesgarani et al. using 75-150 Hz^54^, but we also argue that these cutoffs are still completely arbitrary (an upper limit of 128 or 256 may be more suitable given that many recording technologies are at powers of 2). Regardless, we show results (Fig. S10) in mid gamma band (70-100Hz), which combined with the results of our high gamma band, would include the range 70-128 Hz. There are consistent results across the study. Note that it seems effect sizes become larger at higher Hz, so we suspect that removing the patient with a sampling of 256Hz to include higher frequency bands may still not alter our conclusions.

### GM and WM functional connectivity (FC) differences

To understand GM and WM FC differences, we subdivided the full network (all contacts included) into tissue sub-networks. Four networks in total were studied: (1) the full network (2) GM-only (3) GM-WM, which included only the connections from GM to WM contacts, and (4) WM-only. We analyzed the distribution of FC values by taking the top triangle half of the symmetrical, undirected adjacency matrix, except for the GM-WM tissue sub-network where all FC values were taken (Fig. 2b, 2c, and 4b histograms). We plotted the empirical cumulative density functions (ECDF) of the sub-network FC values (Fig. 2d and 4b). These plots (Fig. 2b-d) show a representative example patient and seizure. Subsequent plots (Fig. 2e, right to 2h) show at the subject-level.

We wanted to answer the biological question if WM connections are higher (or lower, as hypothesized initially) during the ictal state. Continuing our illustration of the example patient plotted in Fig. 2d, we performed a two-sided Kolmogorov-Smirnov test to test the null hypothesis that the two sub-network distributions of FC values are identical during the ictal period. We found evidence against the null hypothesis.

Across all subjects, we took two approaches to visualizing the differences in the distributions of WM and GM FC values. (1) We pooled all FC values within each tissue sub-network across all patients (i.e. all the WM FC values from patient 1 would be pooled with all the WM FC values from patient 2, and so forth for all patients; N = 29). Then (similar to Fig. 2d), we plotted the pooled FC distributions across all patients and the two tissue sub-networks for each state (Fig. 2d, ECDF plots). We see in the ictal state, the WM (blue dotted line) is pulled to the right. (2) In the second visualization approach, we bootstrapped patients and seizures instead of pooling FC values. For *each* re-sampled patient, we found the median FC value for each tissue sub-network and state. We then took the mean across all patients for each tissue sub-network and state. We plotted the means for 10,000 bootstrap simulations for each state in the inset histograms in Fig. 2d. We see that WM FC is lower during the interictal and preictal states, however, WM FC becomes higher during the ictal state.

To directly compare the GM-only and WM-only sub-networks across each patient, we defined Δ*r* as the difference in the median FC value for each patient. Here we had two null hypotheses: (1) Δ*r* during the ictal state (Δ*r*_*ic*_) equals zero, meaning that there is not a difference in the tissue sub-network FC values during the ictal state and (2) Δ*r*_*ic*_ − Δ*r*_*pi*_ equals zero. In other words, rejection of our second null hypothesis would support our alternative hypothesis that the differences in the tissue sub-network FC values become *larger* at the transition to a seizure state.

We show Δ*r* for all patients and states (Fig. 2h, left). To test our two null hypotheses, we performed a Mann-Whitney U rank test for the first hypothesis and a Wilcoxon signed-rank test for the second hypothesis (paired test between preictal and ictal). The means of 1,000 bootstrap simulations of Δ*r* for each state were plotted in Fig. 2h, right.

### Controlling for distances between contacts

Although mid-line GM structures (such as the cingulate gyrus) or more medial structures (such as the limbic system) are targets for implantation, it is reasonable to suspect WM contacts are closer together and may explain the increased FC values. Thus, a covariate in our analyses is distance between contacts. We measured euclidean distance between contacts from the coordinates produced in the electrode localization pipeline above. Although we mitigated the influence of passive volume conduction through bipolar montaging explained above, adjacent contacts may still measure related signals even if montaging methods completely eliminated volume conduction because adjacent neural tissue may be performing similar, independent computation. Contacts closer together have higher FC than contacts further away (Fig. 2e) after fitting a ordinary least squares linear regression line (contact distance is first *log*_10_ transformed).

Here we wanted to test if WM sub-networks still have increasing FC farther from GM tissue (we quantified “farness” using two definitions, distance and depth defined above). We included the distance between contacts as a covariate in both ordinary least squares models using either the WM distance or WM depth as the other covariate (in Python package statsmodel, we used OLS.from_formula: FC value ∼WM definition + contact distance). P-values reported in Fig. 2f show the probability that the covariate of WM distance or depth have no effect on FC values.

### Network Analysis

We took a network analysis approach in studying GM and WM differences. We compared the Full, GM-only, and WM-only networks. We could not feasibly compare the GM-WM networks taking a network analysis approach because these networks are not symmetric (they are not an NxN matrix where N represents the number of nodes in the network). All network measures were calculated using the Brain Connectivity Toolbox for Python, including density, characteristic path length, and transitivity. These network measures were chosen to understand how basic network properties differe between the sub-networks. We also wanted to answer if WM networks are more clustered than GM networks based on the results in Fig. 2 through a measure of transitivity. We performed network analysis on an initial trial of 8 patients (marked in Table 1) and results are shown in Fig. 3. We thresholded the broadband cross correlation functional connectivity matrices at 0.2 and binarized the matrices. After computing the network measures, we compared GM-only and WM-only tissue sub-networks for each state using the Wilcoxon signed-rank test.

### Imaging protocol for structure-function analysis

Prior to electrode implantation, MRI data were collected on a 3T Siemens Magnetom Trio scanner using a 32-channel phased-array head coil. High-resolution anatomical images were acquired using a magnetization prepared rapid gradient echo (MPRAGE) T1-weighted sequence (repetition time = 1810 ms, echo time = 3.51m, flip angle = 9, field of view = 240mm, resolution = 0.94×0.94×1.0 mm3). High Angular Resolution Diffusion Imaging (HARDI) was acquired with a single-shot EPI multi-shell diffusion-weighted imaging (DWI) sequence (116 diffusion sampling directions, b-values of 0, 300, 700, and 2000s/mm2, resolution = 2.5×2.5×2.5 mm3, field of view = 240mm). Following electrode implantation, spiral CT images (Siemens) were obtained clinically for the purposes of electrode localization. Both bone and tissue windows were obtained (120kV, 300mA, axial slice thickness = 1.0mm)

### Diffusion Weighted Imaging (DWI) Preprocessing

HARDI images were subject to preprocessing pipeline QSIPrep to ensure reproducibility and implementation of the best practices for processing of diffusion images^55^. Briefly, QSIPrep performs advanced reconstruction and tractography methods in curated workflows using tools from leading software packages, including FSL, ANTs, and DSI Studio with input data specified in the Brain Imaging Data Structure (BIDS) layout.

### Structural Network Generation

DSI-Studio (http://dsi-studio.labsolver.org, version: March 2021) was used to reconstruct the orientation density functions within each voxel using generalized q-sample imaging with a diffusion sampling length ratio of 1.25^56^. Deterministic whole-brain fiber tracking was performed using an angular threshold of 35 degrees, step size of 1mm, and quantitative anisotropy threshold based on Otsu’s threshold^57^. Tracks with length shorter than 30mm or longer than 800mm were discarded, and a total of 1,000,000 tracts were generated per brain. Deterministic tractography was chosen based upon prior work indicating that deterministic tractography generates fewer false positive connections than probabilistic approaches, and that network-based estimations are substantially less accurate when false positives are introduced into the network compared with false negatives^58^. To calculate structural connectivity between electrode contacts, ROIs of 7mm radius surrounding the contacts were created (Fig. 4a). The ROI volume was selected to equate to the volume of an average ROI in another commonly used neuroimaging atlas, the Automated Anatomical Labeling (AAL) atlas^59–61^. This soft-boundary^46^, contact-specific atlas was used to because we were studying individual signals from each contact and the tissue they recorded from. Structural networks were generated by computing the number of streamlines passing through each pair of ROIs. Streamline counts were log-transformed and normalized to the maximum streamline count, as is common in prior studies^62–65^.

### Structure-Function Correlation

To quantify the relationship between structure and function, we computed the Spearman Rank correlation coefficient between the edges of the structural connectivity networks and the edges of the functional connectivity network (Fig. 4c). Broadband cross correlation was used because it has shown the most robust SFC in epilepsy patients^66^. We divided each structural and functional network in tissue sub-networks: (1) A full network representing equal treatment of GM and WM contacts (2) a GM-only network. (3) and GM-WM network (4) a WM-only network. We correlated each structural and functional sub-network. We re-sampled our data set (see bootstrapping methods below) and compared the the SFC of each sub-network across the four states. We performed the Wilcoxon signed-rank test between the ictal and preictal states for each sub-network to test the following alternative hypothesis: Δ*SFC*_*tissue*_ ≠ 0, where Δ*SFC*_*tissue*_ represents the difference between the ictal and preictal SFC (*SFC*_*ic*_ − *SFC*_*pi*_) for the given tissue sub-network. We hypothesized that Δ*SFC* is higher in any tissue sub-network containing GM because GM is the neural tissue that drives the activity of the brain. Thus WM Δ*SFC* would be lower. We bootstrapped this experiment 1000 times and plot the distribution of the mean Δ*SFC*_*tissue*_ across the re-sampled patients and seizures. We report confidence intervals.

### Outcome Classification

We determined the outcomes of epilepsy surgery using the Engel classification scoring system^67^. We included patients we were ≥ 2 years from surgery to account time where seizure occurrences may relapse. Clinical studies monitoring outcomes are ongoing, however ≥ 2 years was selected because we felt this was ample time to increase the accuracy of the ground-truth effectiveness of the epilepsy surgery. We separated good (Engel I) and poor (Engel II-IV) outcome patients typically done in the literature^35,65,68–73^. Note that we dichotomized good and poor outcomes because we assume that in good outcome patients, the ablated regions contained a well-localized seizure generating area that was correctly targeted. We wanted to understand how this targeted area is related to WM activity. Note that a poor outcome patient does not necessarily mean a “poor” outcome for the patient. Engel III (a poor outcome class in this study) means marked seizure improvement, which greatly enhances qualitty of life in some patients and would be considered a “good” outcome to them. Here we dichotomize Engel I and II-IV, not from a patient point of view, but rather from a neurosurgical point of view in accurately localizing a circumscribed SOZ. In the case of Engel II-IV, patients continue having seizures, and thus the true SOZ was missed (see the discussiion in the main text ‘The Distributed Epilepticc Network Hypothesis”). In this study, poor outcome patients were not recommended for palliative care, meaning their overall clinical findings (including their SEEG recordings) indicated a reasonable suspicion for the target area. Seizure freedom may last for a short time (≤ 2 years). After a unspecified time, brain regions may adapt and seizures may continue. Hence, this is one reason why we included patients with outcomes of follow-up longer than previous studies.

### Functional connectivity of tissue sub-networks in good and poor outcomes

We compared the tissue sub-networks in good and poor outcome patients using methods similar to Fig. 2. We compared the distribution of FC values in (1) the full (2) GM-only (3) GM-WM and (4) WM-only networks between good and poor outcomes during the preictal and ictal states. We tested the null hypothesis that the distribution of FC values in good and poor outcome patients with respect to each sub-network and state are similar. We performed a two-sided Kolmogorov-Smirnov test and outputs are shown in Fig. 5b. We also compared the change in FC (Δ*FC*) from preictal to ictal states (Δ*FC* = *FC*_*ictal*_ − *FC*_*preictal*_) from the contacts contained within the ablation zone to other WM contacts (Fig. 5f). We pooled all FC values from the bootstrapped patients, seizures, and connectivity matrices (see bootstrapping methods below) and plotted the distributions (Fig. 5f, left) similar to Fig. 2g. To compare Δ*FC* on a patient level, we took the median FC value for each patient, and plotted the results in Fig. 5f, right. To test the null hypothesis that good outcome patients have similar FC values from the ablation zone to all WM contacts, we performed the Mann-Whitney U test. To provide more robust statistical methods considering our small sample size of poor outcome patients, re-sampling hypothesis testing was also performed and is explained in detail below.

### Hypothesis testing comparing good and poor outcome patients

We wanted to know if WM activity is different between good and poor outcome epilepsy patients.

We repeated the analysis in Fig. 2h separating good and poor outcome patients with the null and alternative hypotheses defined below:

1. 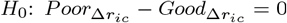. The differences in GM functional connectivity and WM functional connectivity (Δ*r*, differences in *T* issue) is similar for good and poor outcome patients during the ictal period.
2. 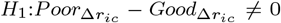. The differences in GM functional connectivity and WM functional connectivity (Δ*r*, differences in *T* issue) is NOT similar for good and poor outcome patients during the ictal period.

After the results from Fig. 5c, we wanted to test the hypothesis that ablated regions in good outcome patients do not increase in correlation to WM regions as much as poor outcome patients during seizures. This hypothesis helps answer the biological question if WM electrodes reveal a distributed, less-localizing nature in the pathophysiology of their epilepsy. Here we formally define the null and alternative hypotheses:

1. 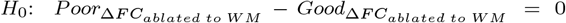. The change in functional connectivity from preictal to ictal states (Δ*FC*) from the ablated GM regions to WM regions is similar for good and poor outcome patients.
2. 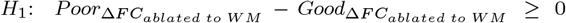. The change in functional connectivity from preictal to ictal states (Δ*FC*) from the ablated GM regions to WM regions is *greater* for poor outcome patients than good outcome patients. Using knowledge and expectations based on prior results, we hypothesize poor outcome patients are greater, therefore, we do a one sided test (see re-sampling hypothesis testing below). We omit the absolute value in eq. 3.

An important note: given the sampling of our data (un-equal seizure number across patients, non-uniform implantation schemes, un-equal tissue sampling, only 17 patients with ≥ 2 years of outcomes reported, etc), we performed both bootstrapping methods and re-sampling hypothesis testing for a robust analysis (see relevant sections below).

### Bootstrapping

The bootstrap is a widely applied and powerful statistical tool^74^. It is useful in our data set where a measure of variability is otherwise difficult to obtain. In our data set, we have a variable amount of seizures per patient. Each patient has a variable implantation strategy with a variable number of recording contacts. Therefore, each patient will also have a different sampling of tissue types with resulting different network sizes. To mitigate these biases, or over-/under-sampling, of any one patient or seizure, we re-sampled with replacement at each level of patient and seizure. For example, we would sample with replacement N patients from our cohort equal to 1x the number of total patients. Then, we would sample with replacement M seizures from each patient equal to the number of seizures that patient has. In this case, because patients are already bootstrapped, seizures can be considered independent. We would bootstrap 10,000 times to ensure patients have equal sampling of seizures.

### Re-sampling hypothesis testing

We employed re-sampling hypothesis testing to understand the differences in WM seizure activity in good and poor outcome patients in Fig. 5. Due to a small sample size of our patients with validated outcome scores ≥ 2 years, we follow the permutation approach to the p-value outlined in James et al., 2021.^74^. As illustrated in their text, this approach is preferable and is considered reliable; we follow Algorithm 13.3 of the text exactly and written here for convenience:

1. Compute the test statistic, *T*, on the original data 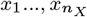, and 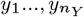.
2. For *b* = 1, …*B*:
  a. Permute the *n*_*X*_ + *n*_*Y*_ observations at random. Call the first *n*_*X*_ permuted observations 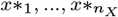, and call the remaining observations 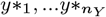.
  b. Compute *T* on the permuted data 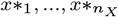 and 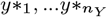, and call the result *T*^∗*b*^
3. The *p*-value is given by eq. 3 below.

We computed the test statistic as the two-sample *t*-statistic Following James et al., 2021.^74^:

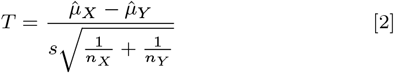

where 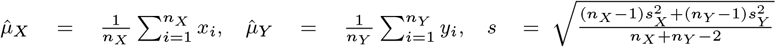 and 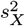 and 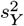 are unbiased estimators of the variances in poor and good outcome patients, respectively. We randomly permute the *n*_*X*_ + *n*_*Y*_ observations B times, and each time we compute eq. 2. We let *T*^∗1^, …,*T*^∗*B*^ denote the values of eq. 2 on the permuted data. These can be viewed as an approximation of the null distribution of *T* under *H*_0_. To compute the *p*-value, we do

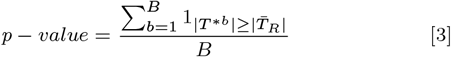

This computes the fraction of permuted datasets for which the value of the test statistic is at least as extreme as the value observed on the original data’s test statistic, *T*. For our re-sampling hypothesis testing approach, we treated each seizure as an independent sample. We also averaged over patients-level and show significant (p<0.05) or trending (p<0.1) results in Fig. S6. Note that we approached the analysis comparing GM and WM throughout the manuscript using multiple methods to ensure we lower our Type II error rates testing the overall question if WM is different from GM and provides unique information about epilepsy.

### Functional connectivity changes in WM as a function of distance from ablation zone between good and poor outcome patients

We measured the change in functional connectivity (Δ*FC*) from preictal to ictal states between all electrode contacts. We determined which contacts recorded neural activity in the suspected seizure onset zone and subsequently ablated. We then determined the FC between the ablated contacts and WM contacts. We measured the distance between those contacts and took the median FC of all the contacts less than or equal to the distance specified on the x-axis of Fig. 2h. Ninety-five percent confidence intervals are plotted. The results shown is the outcome of one bootstrap simulation (see bootstrapping methods above).

A note on the ablation zone: In good outcome patients, we assume that the ablation zone contained the “true” seizure onset zone, or alternatively, contained a seizure area that was well localized to that region. In poor outcome patients, the ablated regions were targeted because it followed the above assumptions in good outcome patients, but the outcomes would not be known yet. It was the suspected SOZ. However, our results show that in poor outcome patients, this suspected SOZ becomes more correlated to distant WM regions, pointing to a more distributed, non-localizing nature of their seizure pathophysiology even in the face of all clinical evidence pointing to otherwise (or else they would not have been recommended epilepsy surgery, but rather alternative treatment options such an implantable neurostimulating devices).

### Data availability and Reproducibility

All code files used in this manuscript are available at https://github.com/andyrevell/revellLab. All de-identified raw and processed data (except for patient MRI imaging) are available for download on Box. Link provided on GitHub. The GitHub repository used to analyze the data is also contained within Box. Raw imaging data is available upon reasonable request from Principal Investigator K.A.D.; tractography files generated from the imaging data are readily available on Box. iEEG snippets used specifically in this manuscript are contained within the Box data folder, while full iEEG recordings are publicly available at https://www.ieeg.org. The Python environment for the exact packages and versions used in this study in contained in the environment directory within the GitHub. The QSIPrep docker container was used for DWI preprocessing.

#### Acknowledgements

We thank Adam Gibson, Carolyn Wilkinson, Jacqueline Boccanfuso, Magda Wernovsky, Ryan Archer, Kelly Oechsel, members of Andrew’s Thesis Committee, Braden Kelner (for editing), and all other members and staff of the Center for Neuroengineering and Therapeutics for their continued help and support in this work.

## Supplementary Materials

Please see supplemental materials below.

### Glossary

1. **Ablated contacts**. Contacts which fell in the (eventually) ablated region
2. **GM**: Gray Matter. See tissue definitions in the methods section for details on how contacts were classified as either GM or WM.
3. **WM**: White Matter. See tissue definitions in the methods section for details on how contacts were classified as either GM or WM.
4. **GM contact**. A contact defined as GM, using tissue definitions detailed in the methods section.
5. **WM contact**. A contact defined as WM, using tissue definitions detailed in the methods section.
6. Δ **SFC**: The change in SFC between ictal and preictal stats (*SFC*_*ictal*_ − *SFC*_*preictal*_). This indicates whether or not the change in functional connectivity is congruent with the underlying structural connectivity.
7. Δ **FC**. The change in functional connectivity between ictal and preictal states. *FC*_*ictal*_ − *FC*_*preictal*_. This measures if two contacts are increasing or decreasing in the functional connectivity values when a seizure evolves.
8. **ii, pi, ic, po**. Interictal, preictal, ictal, postictal states.
9. **T1w**. T1-weighted MRI image.
10. **Contact**. A single sensor on an electrode that records LFP. Not to be confused with the electrode itself (an entire “stick” of an SEEG)
11. **Electrode**. Not to be confused with contact.
12. **SEEG**: Stereoelectroeenccephalography.
13. **SNR**: Signal-to-noise ratio. See methods section.
14. **ECoG**: Electrocorticography.
15. **Functional connectivity (FC)**. The statistical relationship between two signals (two contacts in this study). Many FC measurements exists, including but not limited to Pearson correlation, Spearman Rank correlation, Cross correlation, and Coherence. We primarily use Cross correlation in this manuscript.
16. **Structural connectivity (SC)**. The physical relationship between two brain regions. We use streamline counts in this manuscript from High Angular Resolution Diffusion Imaging.
17. **Bootstrapping**. A robust statistical tool. See methods section.
18. **Re-sampling hypothesis testing**. A robust statistical analysis technique for testing hypothesis using permutation approaches. See methods section.

**Fig. S1.**
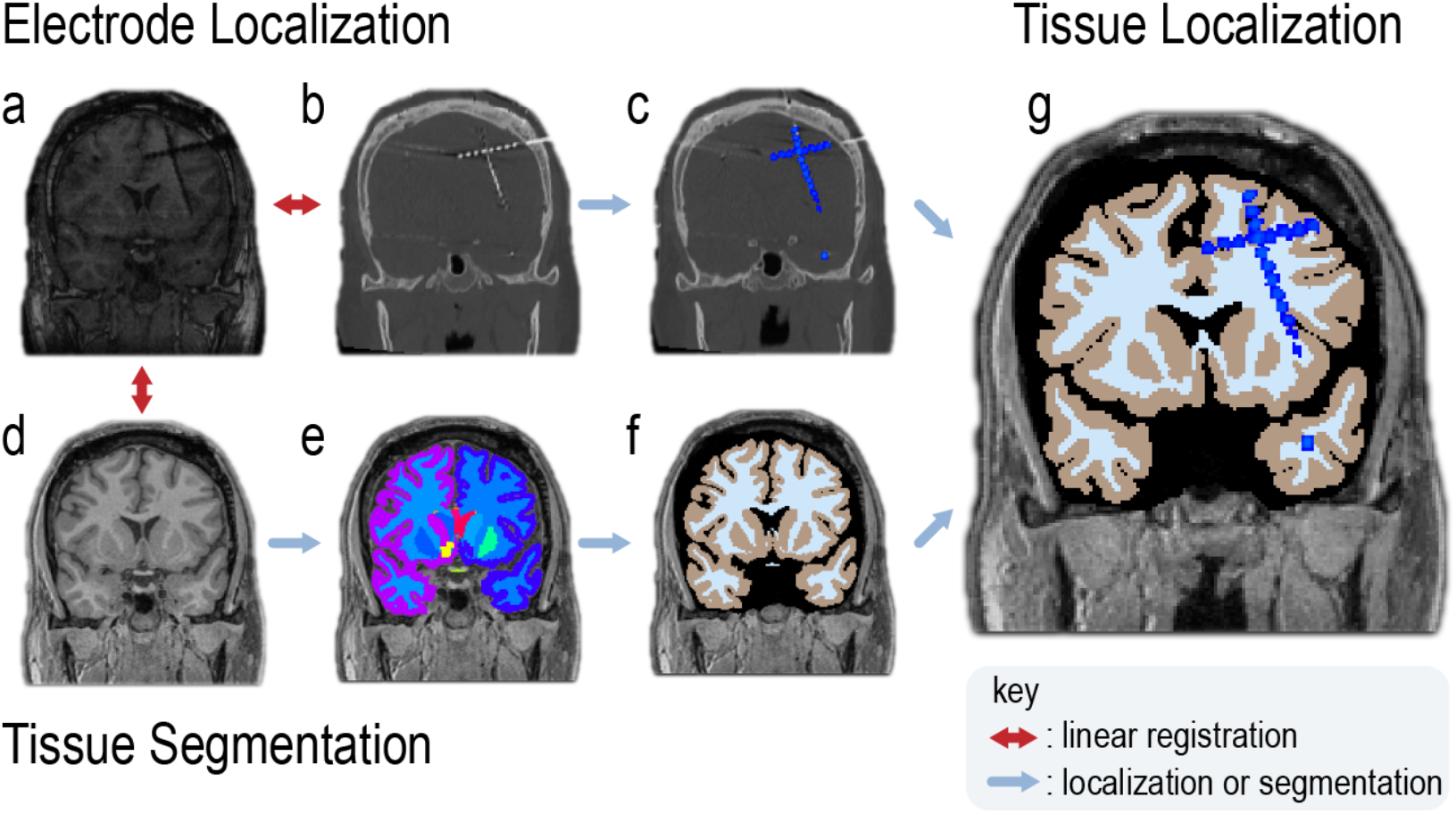
Overview of Localization and Segmentation. **a**, Implant T1w MRI. **b**, Implant CT. **c**, Implant CT SEEG contacts are segmented and localized in native coordinates. **d**, Pre-implant 3T T1w MPRAGE. Usually higher quality imaging (1mm isotropic) was used for tissue segmentation. If patient did not undergo high resolution imaging, then a clinical T1w pre-implant image is used. **e**, Segmentation output by Freesurfer showing tissue parcellations. **f**, Conversion of tissue parcellation into labels of GM, WM, CSF, or outside of the brain. **g**, Combination of entire pipeline to localize electrode contacts to tissue. Images (a) and (b) are linear registered using FSL’s flirt command so that contact coordinates localized in native CT space can be converted to image (a)’s space. Images (a) and (d) are linear registered so that contact coordinates in image (a)’s space can be converted to image (d)’s space. Once contacts are localized to image (d)’s space and image (d) has been segmented, coordinates and segmentation are combined to produce image (g). WM classification is determined using two definitions shown in Fig. S2.

**Fig. S2.**
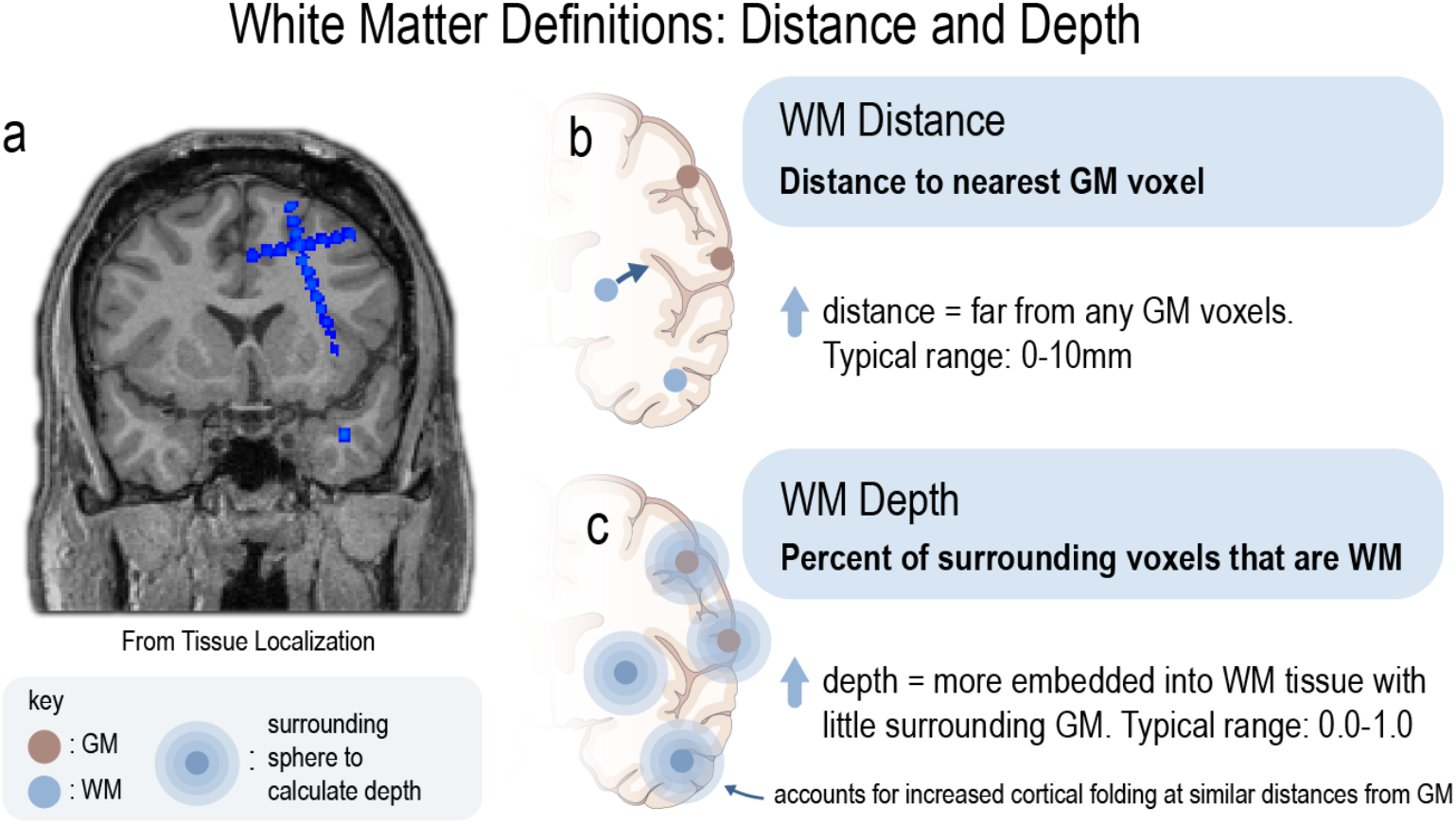
White Matter Definitions: Distance and Depth. **a**, Contacts are localized to tissue. See Fig. S1. **b**, Binary classification of WM and GM contacts can be found by measuring the distance of the contact’s center to the nearest GM voxel. **c**, Contacts can be classified by WM depth. This is the percent of surrounding voxels (within a 1cm diameter sphere) that are WM. This is closely related to the PTD measurement in Mercier et al. 2017^7^ (see Methods). A benefit of WM depth is that it accounts for cortical folding and small errors in tissue segmentation. WM contacts near the cortical surface, but a few voxels from GM will have low WM depth because they are still surrounded by brain sulci, and thus many GM voxels. Furthermore, a single voxel mis-segmented as GM can affect WM distance. Therefore, we primarily used WM depth.

**Fig. S3.**
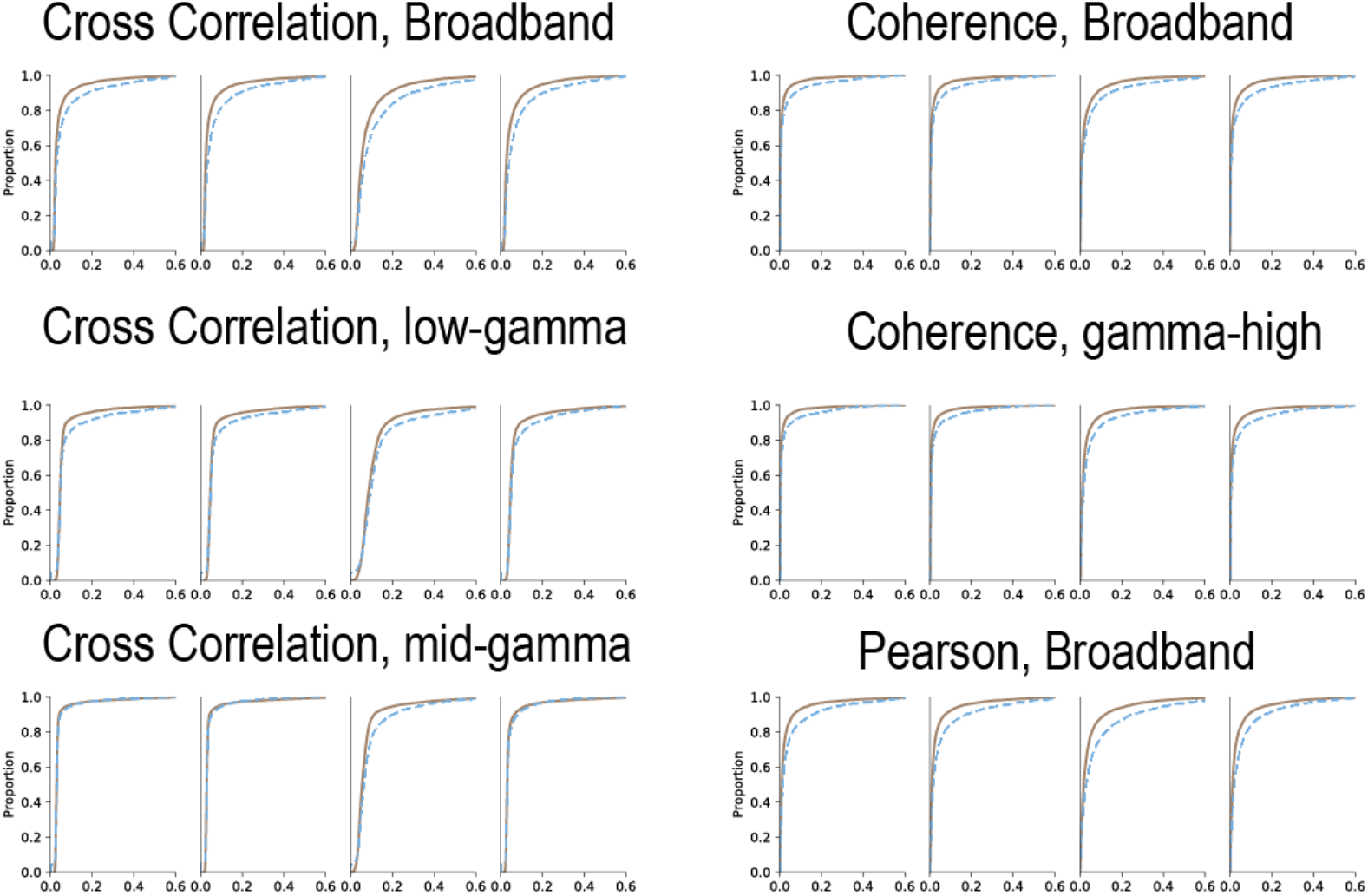
GM and WM distribution of functional connectivity value: Select frequency bands and other functional connectivity measurements. We repeat analysis from Fig. 2g using select frequency bands and functional connectivity measurements. For all frequency bands and functional connectivity measurements, repeating Fig. 2h, see **Fig. S4**. We focus on high-*γ*-band and cross correlation in the main text as detailed previously. Our central hypotheses related to seizure activity, therefore, we tested our findings using high-*γ*. However, we note more striking differences in other frequency bands and functional connectivity measurements. We provide information here and in **Fig. S4** to aid research using this manuscript for a basis.

**Fig. S4.**
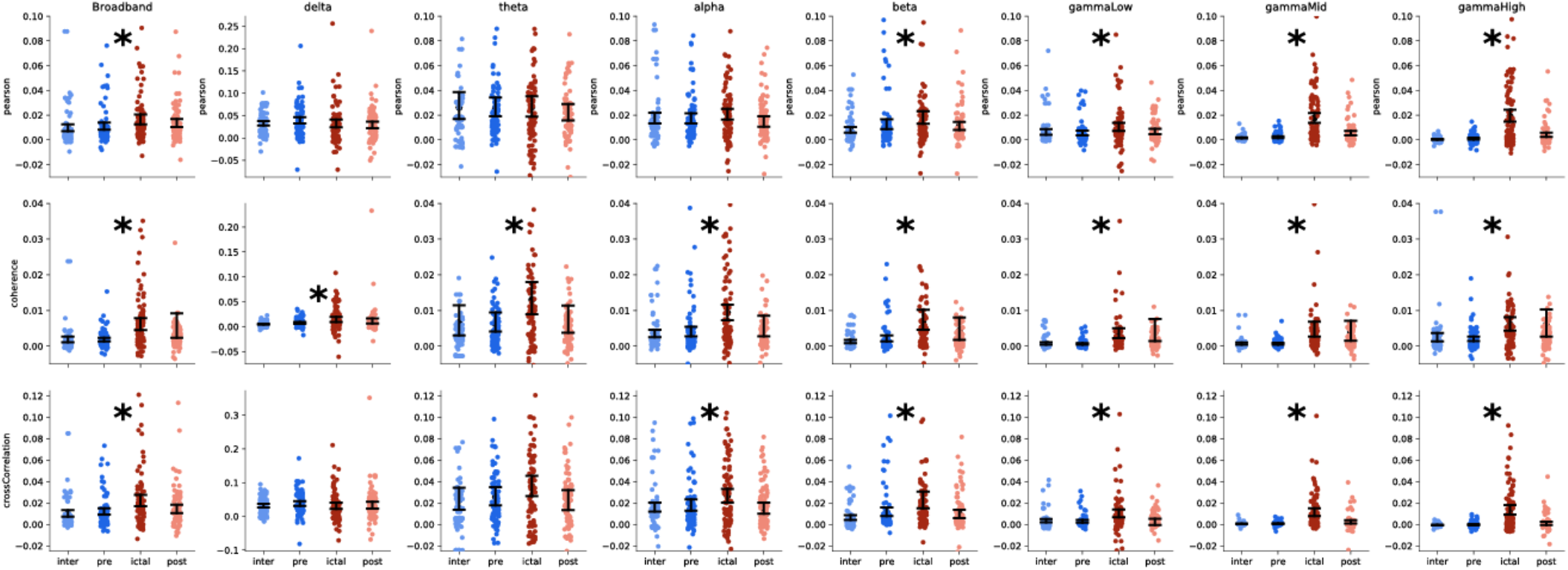
Differences in GM and WM (Δ*r*) for all frequency bands and functional connectivity measurements. We repeat analysis from Fig. 2h showing all frequency bands and functional connectivity measurements. Top row: Pearson correlation. Middle row: Coherence. Bottom row: Cross Correlation. Left-to-right: Broadband, delta, theta, alpha, beta, low-gamma, mid-gamma, high-gamma. See methods on frequency band definitions. We note results may be biased based on the choice of frequency bands, therefore, we provide a more comprehensive analysis here. We selected high-*γ*-band and cross correlation prior to conduction of the study based on prior knowledge about epilepsy and functional connectivity measurements from the literature. Error bars are 95% confidence intervals. Asterisk represent significant findings between the statistical comparison of ictal and preictal Δ*r*. Multiple comparison correction is not performed because we do not use these results subsequently in the manuscript, but we provide the analysis here for reference in other studies.

**Fig. S5.**
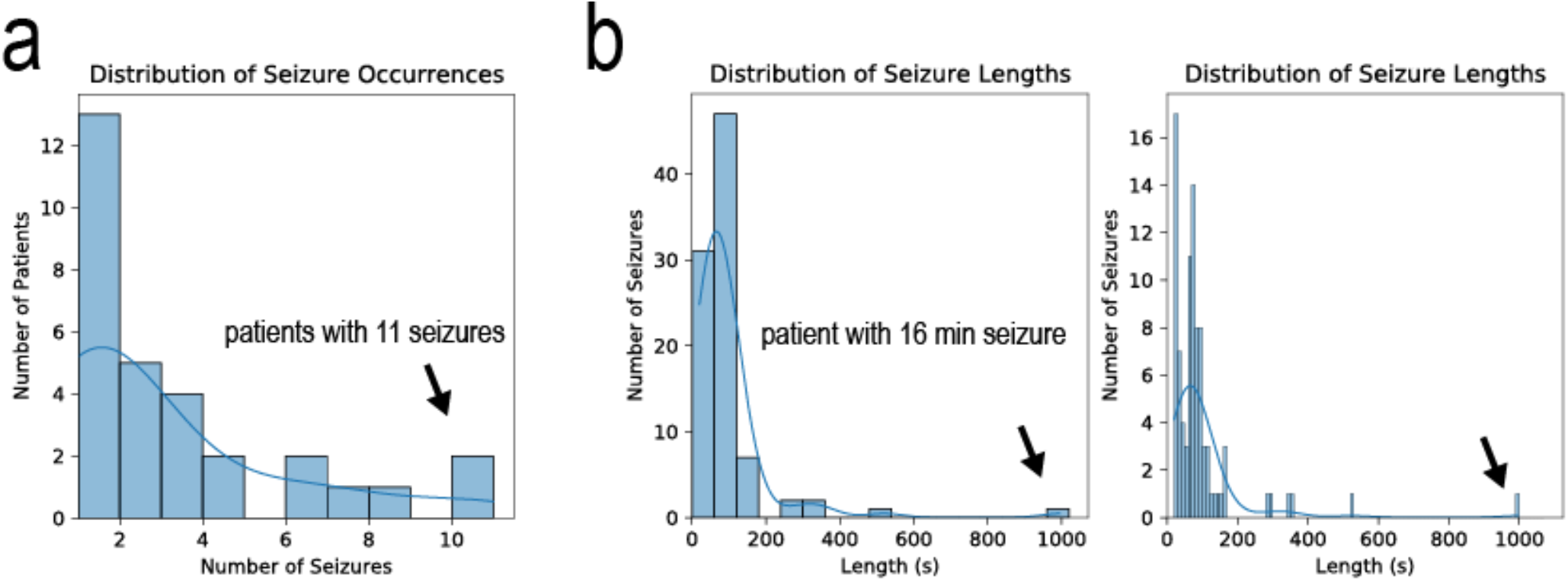
Distribution of seizure occurrences and lengths. **a**, The distribution of seizures for each patient. Most patients (N=13) had one seizure annotated. Two patients had 11 seizures annotated. **b**, The distribution of seizure lengths. Bins of 60 seconds are shown on the left, and bins of 10 seconds are shown on the right. One seizure was approximately 16 minutes long while most seizures were between 1 and 2 minutes long.

**Fig. S6.**
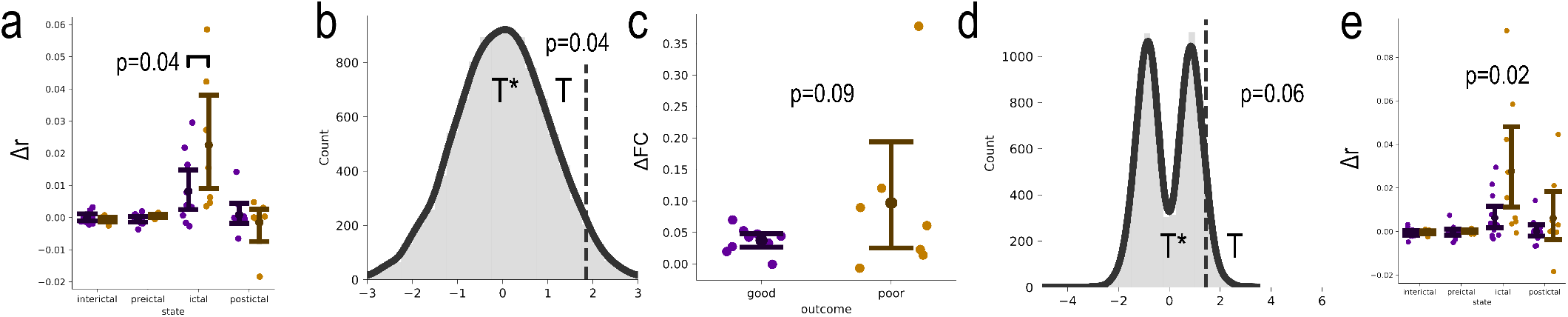
Repeat analysis of Fig. 5 on a per-patient level. **a**, Repeat of Fig. 5c. **b**, Repeat of Fig. 5d, top. **c**, Repeat of Fig. 5f, right. **d**, Repeat of Fig. 5g, top. **e**, Repeat of Fig. S6a with all patients having an Engel outcome score reported, not just ≥2 years (See Table. S1. An important point to stress is that the Distributed Epileptic Hypothesis described in the main text may apply too only *some* poor outcome patients. The reasoning is that 1) the SOZ is still a focal region, but missed during ablation surgery, or (2) the SOZ appears focal to clinicians, but the underlying pathophysiology in some patients points to a more336distributed interaction of brain regions and interconnecting WM pathways^29^. The heterogeneity of poor outcome patients itself proves problematic when poor outcome patients are less common that good outcome patients across epilepsy centers. We show low variation in WM measurements in good outcome patients, but high variability in poor outcome patients. This indicates that different pathophysiological processes are going on in poor outcome patients, however, similar pathophysiological processes are going on in good outcome patients. WM recordings gives better clues to the underlying pathophysiology of epilepsy and thus more research needs to b done. We provide a seizure-level analysis in the main text rather than on a patient-level shown here to understand the variation amongst all seizures. A study with patients from a combination of many epilepsy centers will need to be conducted to improve our Type II error rate. Here we show significance (p<0.05) and trends (p<0.1(in our findings with significance definite ed as an arbitrary cutoff of p<0.05. One-sided independent t-test are performed. We caution to base future work and hypotheses on a single plot in this manuscript, however, we provide ample evidence from multiple analysis approaches to robustly claim that GM and WM have differences related to epilepsy pathophysiology, and that WM recordings should not be ignored.

**Fig. S7.**
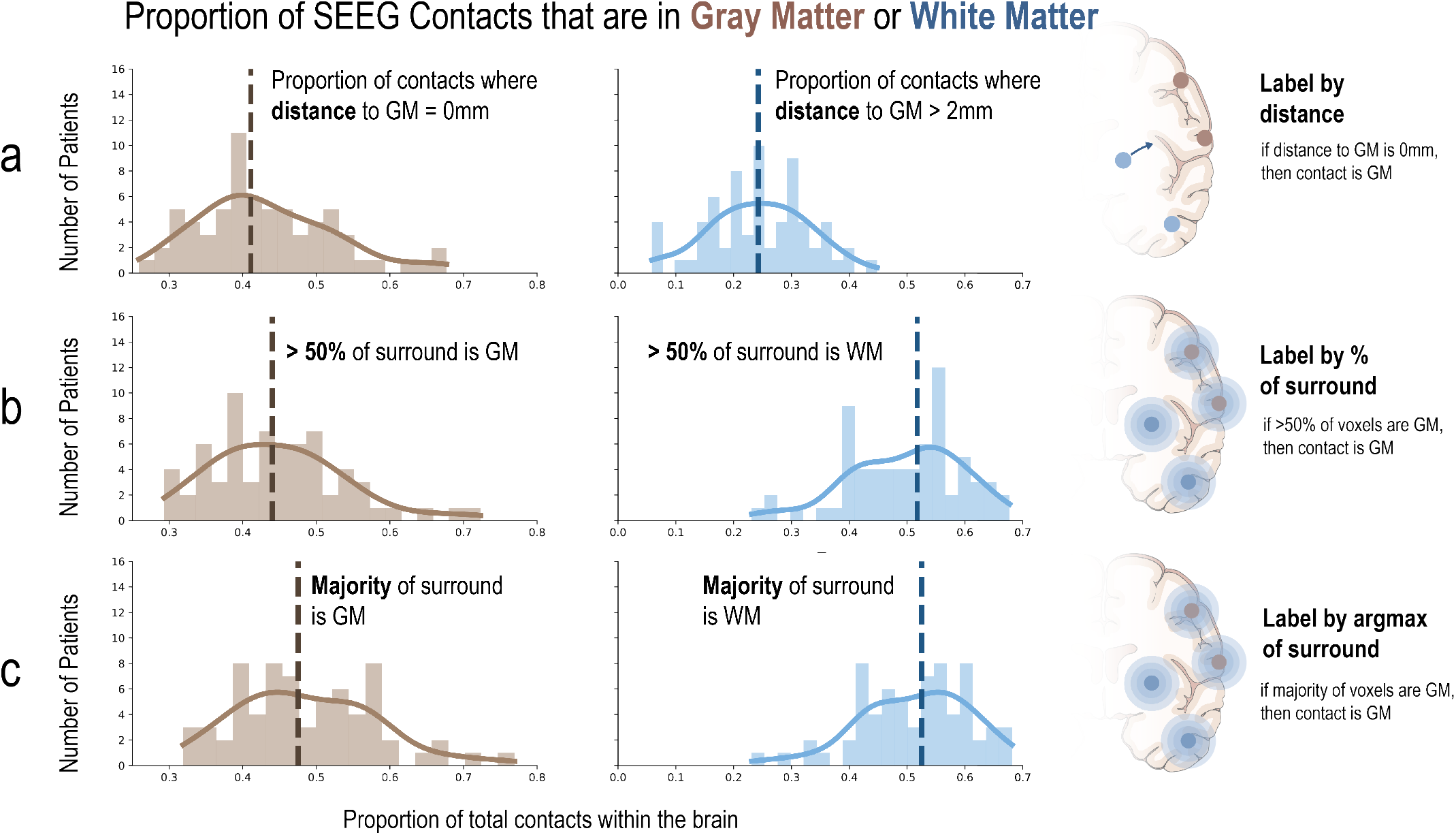
Proportion of contacts in either gray matter (GM) or white matter (WM) **a**, We analyzed 68 SEEG patients. Out of all electrodes that fell within the brain parenchyma (either gray matter or white matter), 41% of contacts fell in a gray matter voxel (dashed vertical line) and 24% of contacts fell in a white matter voxel but also >2mm away from gray matter. This definition reflects one of the definitions used in the main text. Dashed vertical lines represents the median of all patients. **b**, To account for the nuances of any one specific definition, we used two additional approaches to labeling a contact. We analyzed all voxels surrounding the contact centroid within a 1cm diameter and tabulated the number of voxels that were gray matter, white matter, CSF, or outside the brain. A contact was defined as gray matter if >50% of voxels fell in gray matter. If a contact did not have either >50% gray matter or >50% white matter voxels, then it was not defined as either. We report that 44% of contacts were in gray matter and 51% of contacts were in white matter using this definition. **c**, We also labeled each contact using the majority of surrounding voxels (the argmax, rather than >50%). The median patient had 48% of contacts with a majority of gray matter surrounding voxels and 52% of contacts with a majority of white matter surrounding voxels. Panels B and C reflect that approximately half of SEEG electrode contacts fall in white matter.

**Fig. S8.**
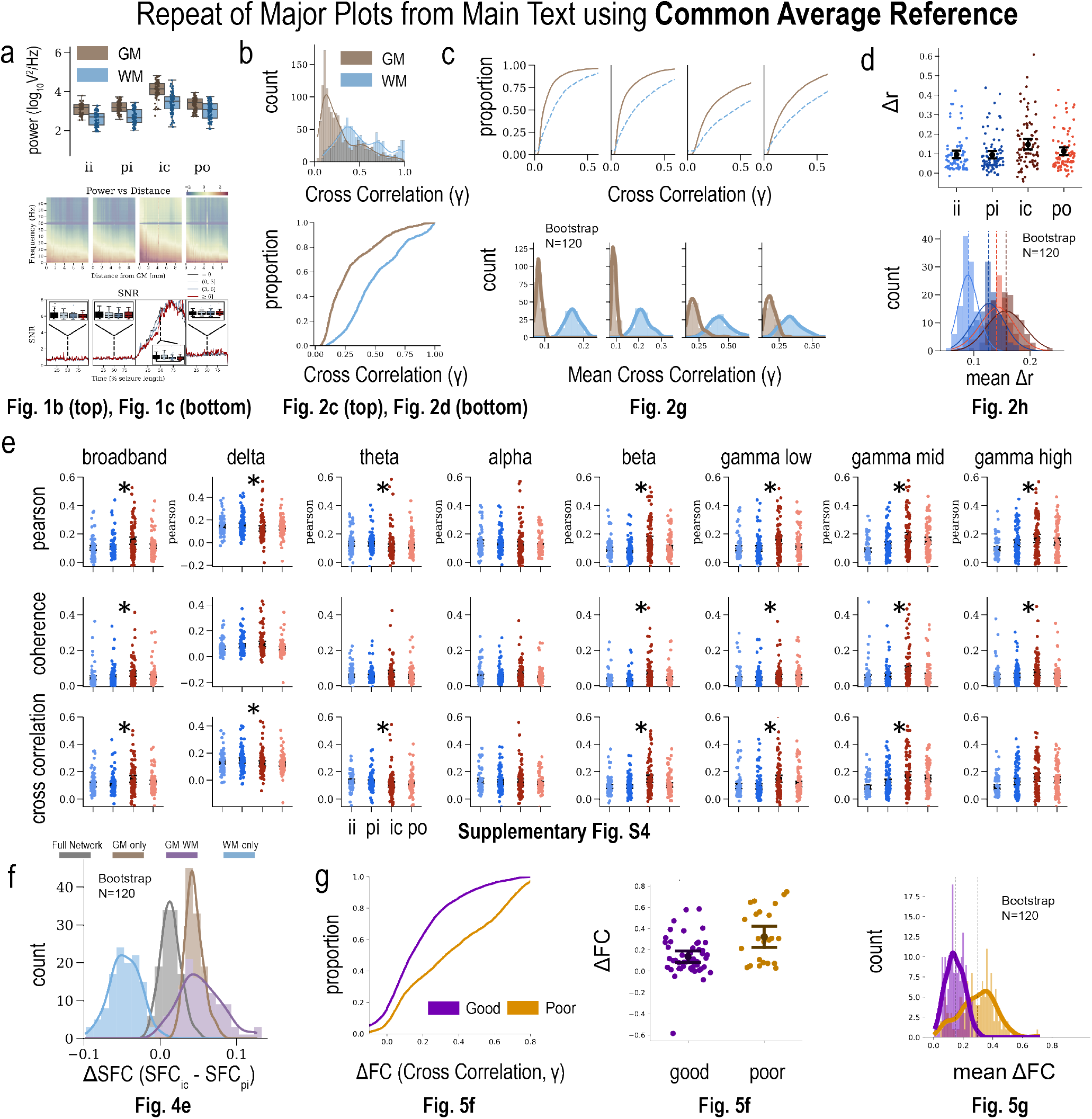
Common Average Referencing (CAR) shows similar results across the entire study as bipolar montage. **a**, Repeat of Fig. 1b (top) and Fig. 1c (bottom) showing similar results with bipolar montage. **b**, Repeat of Fig. 2c (top) and Fig. 2d (bottom) showing similar results with bipolar montage. These plots show results on an individual level from an example patient (the same patient and seizure shown in Fig. 2c and Fig. 2d). **c**, Repeat of Fig. 2g showing similar results with bipolar montage for all patients combined. WM-WM correlations are higher than GM-GM correlations, particularly in the ictal phase. **d**, Repeat of Fig. 2h showing similar results with bipolar montage. **e**, Repeat of Supplementary Fig. S4 showing similar results with bipolar montage across multiple frequency bands. **f**, Structure-Function Correlation. Repeat of Fig. 4e. Correlations between WM signals are not related to their underlying structural connectivity, however, correlations of WM to other GM areas are related to their underlying structural connectivity (similar results with bipolar montage). **g**, Good vs. Poor outcome ablation analysis. Repeat of Fig. 5f (left and middle) showing similar results with bipolar montage. Repeat of Fig. 5g. (right) showing correlations between the ablation site and other WM signals are higher for poor outcome patients (similar results with bipolar montage). In cases where we bootstrapped, we performed 120 permutations (12 cores × 10 loops) rather than 10,000 permutations here for computational efficiency.

**Fig. S9.**
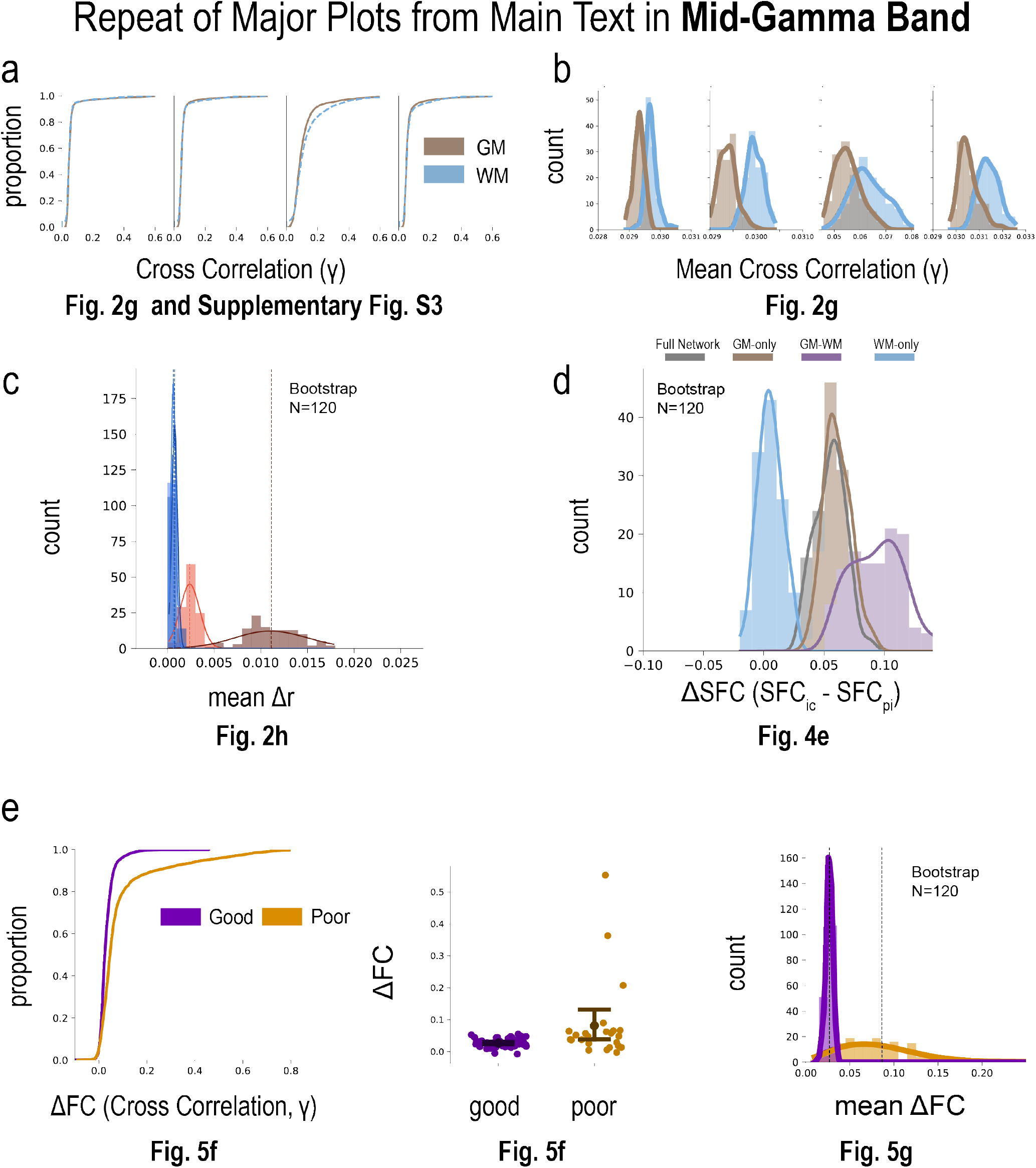
Mid-gamma band shows similar results across the entire study as high gamma. **a**, Repeat of Fig. 2g showing similar results with mid gamma as high-gamma. **b**, Repeat of Fig. 2g showing similar results with mid gamma as high-gamma. **c**, Repeat of Fig. 2h showing similar results with mid gamma as high-gamma. **d**, Structure-Function Correlation. Repeat of Fig. 4e showing similar results with mid gamma as high-gamma. Correlations between WM signals are not related to their underlying structural connectivity, however, correlations of WM to other GM areas are related to their underlying structural connectivity. **e**, Good vs. Poor outcome ablation analysis. Repeat of Fig. 4f (left and middle) and Fig. 4g (right) showing similar results with mid gamma as high-gamma. Correlations between the ablation site and other WM signals are higher for poor outcome patients.

**Fig. S10.**
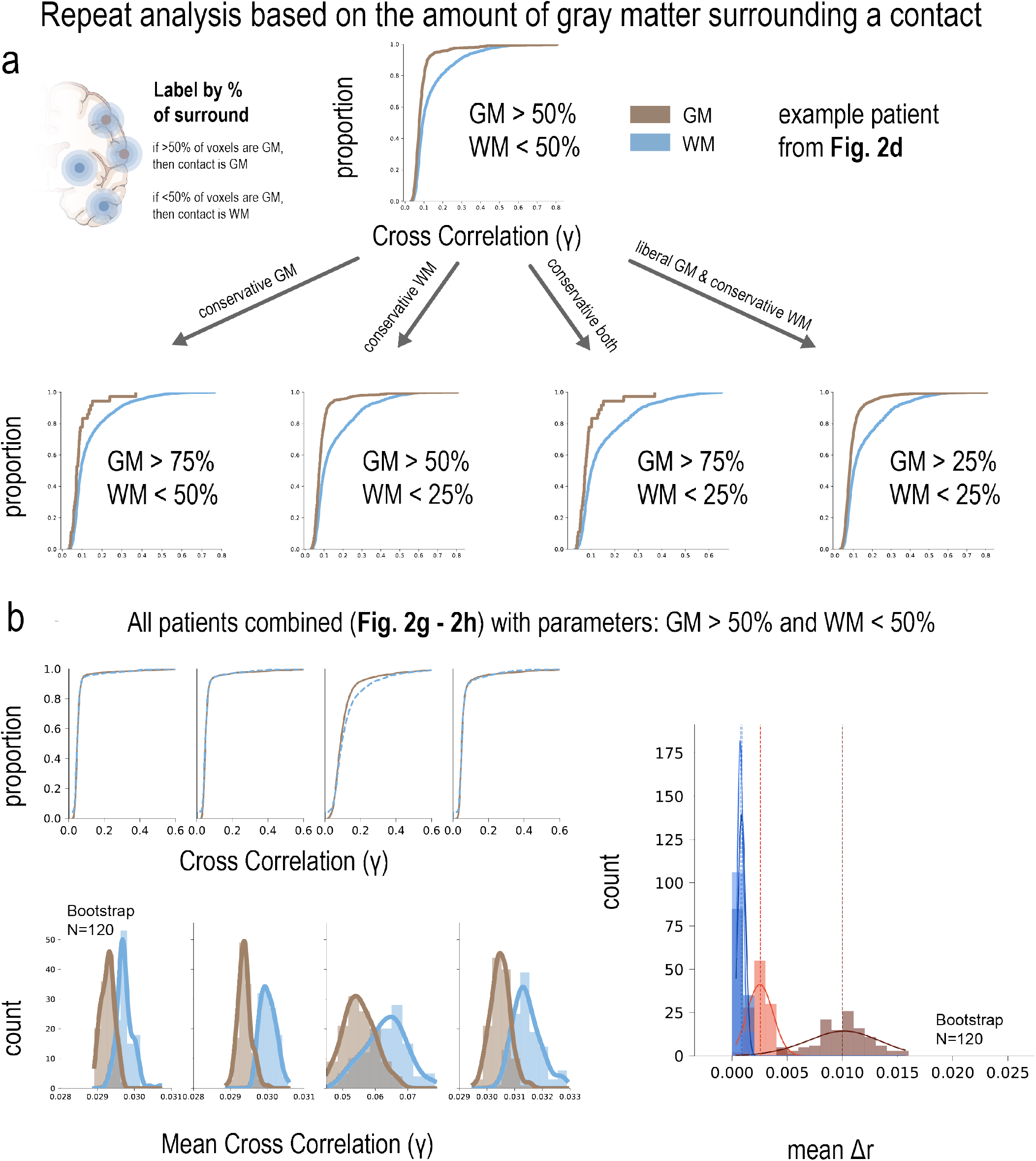
Repeat analysis showing similar results with tissue definition stratified by the amount of gray matter surrounding a contact. **a**, A gray matter contact in this instance is defined by >50% of the surrounding voxels as gray matter, and a white matter contact is <50% of the surrounding voxels as gray matter. As stated in the main text, we have already shown similar results across different tissue definitions and parameters. The results here further support that the nuances of our parameters (including montage selection, frequency bands, functional connectivity definitions) would not change the overall conclusions of our manuscript.

**Table S1.**
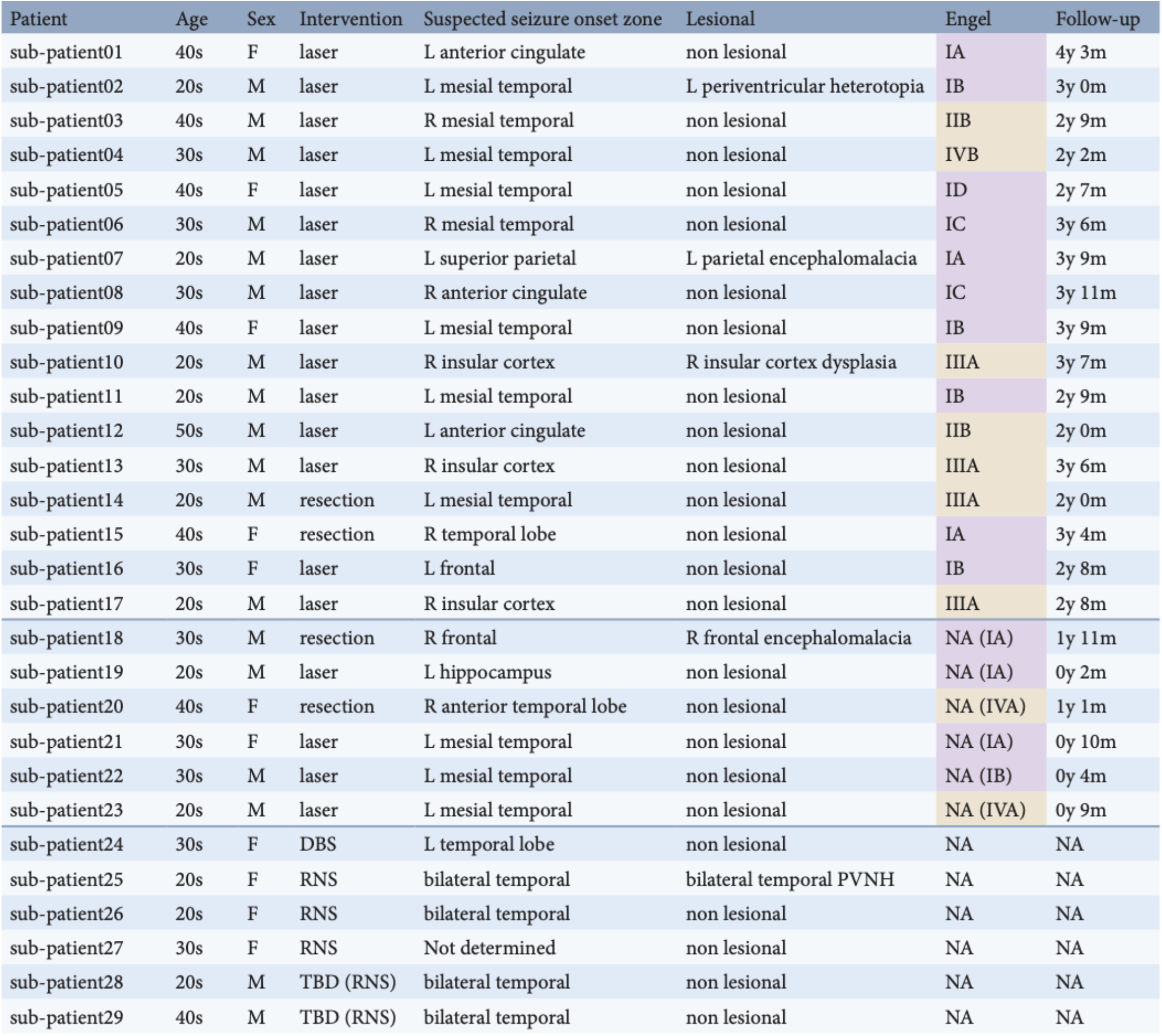
Patient table. We include patient demographics, suspected seizure onset zone (SOZ), lesional status, reported Engel outcome score, and the follow-up time between the patient’s intervention and medical chart review used to determine the outcome score. Note, we masked identifying information as much as possible by reporting approximate age (e.g. “40s”). For example, sub-patient07 is a male in their 20s who underwent laser ablation of their left superior parietal encephalomalacia (positive lesional status), who is reported as a good outcome upon 3 years and 9 months of follow-up after the ablation. Also note that we did not use patients with less than 2 years of follow-up time (as originally planned before conducting the analysis), and we denote the Engel outcome scores of these patients with “NA”. However, we do include what their Engel outcome would be reported from their most recent follow-up. The inclusion of all patients, regardless of follow-up time, yields larger differences between good and poor outcome patients (see Fig. S6e for an analysis including all patients). Only two patients with Engel outcome scores underwent resection rather than ablation, however, in the main text we generalize to writing about ablation only despite a very small minority of patients underwent resection. **M**, Male; **F**, Female; **RID**, Research ID number; **DBS**, Deep brain stimulation; **L**, left; **R**, Right; **NA**, Not Applicable ; **TBD (RNS)**, To be determined, but recommended RNS

| Patient | Age | Sex | Intervention | Suspected seizure onset zone | Lesional | Engel | Follow-up |
| --- | --- | --- | --- | --- | --- | --- | --- |
| sub-patient01 | 40s | F | laser | L anterior cingulate | non lesional | IA | 4y 3m |
| sub-patient02 | 20s | M | laser | L mesial temporal | L periventricular heterotopia | IB | 3y 0m |
| sub-patient03 | 40s | M | laser | R mesial temporal | non lesional | IIB | 2y 9m |
| sub-patient04 | 30s | M | laser | L mesial temporal | non lesional | IVB | 2y 2m |
| sub-patient05 | 40s | F | laser | L mesial temporal | non lesional | ID | 2y 7m |
| sub-patient06 | 30s | M | laser | R mesial temporal | non lesional | IC | 3y 6m |
| sub-patient07 | 20s | M | laser | L superior parietal | L parietal encephalomalacia | IA | 3y 9m |
| sub-patient08 | 30s | M | laser | R anterior cingulate | non lesional | IC | 3y 11m |
| sub-patient09 | 40s | F | laser | L mesial temporal | non lesional | IB | 3y 9m |
| sub-patient10 | 20s | M | laser | R insular cortex | R insular cortex dysplasia | IIIA | 3y 7m |
| sub-patient11 | 20s | M | laser | L mesial temporal | non lesional | IB | 2y 9m |
| sub-patient12 | 50s | M | laser | L anterior cingulate | non lesional | IIB | 2y 0m |
| sub-patient13 | 30s | M | laser | R insular cortex | non lesional | IIIA | 3y 6m |
| sub-patient14 | 20s | M | resection | L mesial temporal | non lesional | IIIA | 2y 0m |
| sub-patient15 | 40s | F | resection | R temporal lobe | non lesional | IA | 3y 4m |
| sub-patient16 | 30s | F | laser | L frontal | non lesional | IB | 2y 8m |
| sub-patient17 | 20s | M | laser | R insular cortex | non lesional | IIIA | 2y 8m |
| sub-patient18 | 30s | M | resection | R frontal | R frontal encephalomalacia | NA (IA) | 1y 11m |
| sub-patient19 | 20s | M | laser | L hippocampus | non lesional | NA (IA) | 0y 2m |
| sub-patient20 | 40s | F | resection | R anterior temporal lobe | non lesional | NA (IVA) | 1y 1m |
| sub-patient21 | 30s | F | laser | L mesial temporal | non lesional | NA (IA) | 0y 10m |
| sub-patient22 | 30s | M | laser | L mesial temporal | non lesional | NA (IB) | 0y 4m |
| sub-patient23 | 20s | M | laser | L mesial temporal | non lesional | NA (IVA) | 0y 9m |
| sub-patient24 | 30s | F | DBS | L temporal lobe | non lesional | NA | NA |
| sub-patient25 | 20s | F | RNS | bilateral temporal | bilateral temporal PVNH | NA | NA |
| sub-patient26 | 20s | F | RNS | bilateral temporal | non lesional | NA | NA |
| sub-patient27 | 30s | F | RNS | Not determined | non lesional | NA | NA |
| sub-patient28 | 20s | M | TBD (RNS) | bilateral temporal | non lesional | NA | NA |
| sub-patient29 | 40s | M | TBD (RNS) | bilateral temporal | non lesional | NA | NA |

